# Reproducible Neuronal Components found using Group Independent Component Analysis in Resting State Electroencephalographic Data

**DOI:** 10.1101/2023.11.14.566952

**Authors:** John Fredy Ochoa-Gómez, Yorguin-José Mantilla-Ramos, Verónica Henao Isaza, Carlos Andrés Tobón, Francisco Lopera, David Aguillón, Jazmín Ximena Suárez

**Affiliations:** Grupo Neuropsicología y Conducta (GRUNECO), Facultad de Medicina, Universidad de Antioquia, Calle 67 N 53 108, Medellín, 050001, Antioquia, Colombia.; Grupo de Neurociencias de Antioquia (GNA), Facultad de Medicina, Universidad de Antioquia, Calle 67 N 53 108, Medellín, Colombia, Medellín, 050001, Antioquia, Colombia.; Semillero de Investigación Neurociencias Computacionales (NeuroCo), Facultad de Medicina & Facultad de Ingeniería, Universidad de Antioquia, Calle 67 N 53 108, Medellín, Colombia, Medellín, 050001, Antioquia, Colombia.

**Keywords:** EEG, Multi-site, Reproducible, group, ICA, Neural, Components

## Abstract

**Objective:** Evaluate the reliability of neural components obtained from the application of the group ICA (gICA) methodology to resting-state EEG datasets acquired from multiple sites.

**Methods:** Five databases from three sites, covering a total of 292 healthy subjects, were analyzed. Each dataset was segmented into groups of 15 subjects, for a total of 19 groups. Data were pre-processed using an automatic pipeline leveraging robust average referencing, wavelet-ICA and automatic rejection of epochs. On each group, stable gICA decompositions were calculated using the ICASSO methodology through a range of orders of decompositions. Each order was characterized by reliability and neuralness metrics, which were evaluated to select a single order of decomposition. Finally, using the decompositions of the selected order, a clustering analysis was performed to find the common components across the 19 groups. Each cluster was characterized by the mean scalp map, its dipole generator with its localization in Talairach coordinates, the spectral behavior of the associated time-series of the components, the assigned ICLabel class and metrics that reflected their reproducibility.

**Results:** Lower order of decompositions benefits the gICA methodology. At this, using an order of ten, the number of reproducible components with high neuronal information tends to be around nine. Of these, the bilateral motor, frontal medial, and occipital neuronal components were the most reproducible across the different datasets, appearing in more than 89% of the 19 groups evaluated.

**Conclusion:** We developed a workflow that allows finding reproducible spatial filters between different data sets. This contributes to the improvement of the spatial resolution of the EEG as a brain mapping technique.

## 1 Introduction

Electroencephalography (EEG) is a portable, noninvasive, and low-cost alternative to visualize brain function. It benefits from a high temporal resolution that is closer to the timescale of neuronal dynamics (da Silva, 2013). The interest in using EEG signals to understand cognitive processes has been recurrent during the last 90 years. The strategy followed by most researchers has been essentially correlative. In most studies, a specific task is designed to execute while simultaneous EEG signals are recorded. Thereafter, researchers extract quantitative indexes, and a statistical approach is used to find significant associations between cognitive scales and the EEG features (Schomer, Lopes da Silva, Michel, & He, 2017).

EEG is characterized by its oscillatory behavior. Oscillations occur at different frequencies from infra-slow (0.5 Hz) to very fast (100 Hz or even more). EEG signals are described in terms of frequency bands with limits defined without any relation to neurophysiological processes. Statistical factor analysis of EEG spectral values yields clusters with considerable overlap with classically accepted frequency bands: infra-slow (*<* 0.2 Hz), *δ* ([0.2, 3.5] Hz), *θ* ([4, 7.5] Hz), *α* ([8, 13] Hz), *β* ([14, 30] Hz), *γ* ([30, 90] Hz) and High Frequency Oscillations (HFO; *>* 90Hz). The oscillations at the lower end of the spectrum tend to engage larger spatial domains, whereas the oscillations at higher frequencies are restricted to specific cortical areas (da Silva, 2013).

The EEG reflects the mass action of neural networks from different brain systems, and thus provides a direct, integrative and noninvasive view of human brain function. The major generators of EEG are extended patches of gray matter, polarized through synchronous synaptic inputs (Buzsáki, Anastassiou, & Koch, 2012). In the cortex, the patches contain thousands of cortical columns, with large pyramidal cells aligned perpendicularly to cortical surfaces (Brandeis, Michel, & Amzica, 2009). The pyramidal cells are the mains, but not sole, contributors to EEG generation. Thus, scalp potentials are the weighted sum of all active currents in the brain that generate open fields. The sum of all open field generators forms an equivalent dipole generator (Brandeis et al., 2009). The concept of dipole generator is not only useful to understand the scalp EEG generation, but also to comprehend the Independent Component Analysis (Delorme, Palmer, Onton, Oostenveld, & Makeig, 2012).

The utility of EEG as a functional neuroimaging method is underestimated, being the EEG commonly associated with a visual interpretation by a skilled neurophysiologist. On the other hand, neuroimaging has been most related to Magnetoencephalography which is a technique that measures similar activity with similar limitations (Schomer et al., 2017). The main concern for brain mapping techniques is the capacity to directly relate the electrical activity measured in an electrode, to the brain zone under that electrode.

By electrical conductance, far-field potentials generated by the cortex sum linearly at every scalp electrode. Thus, EEG data recorded at a single electrode are a weighted linear mixture of underlying cortical sources signals. This spatial mixing by volume conduction is the reason why EEG has been designated as a technique of poor spatial resolution (Onton & Makeig, 2006). The Independent Component Analysis (ICA) decomposes the multichannel EEG into maximally independent processes related to brain activity or artifacts, cancels the mixing process, and opens the possibility to infer the origin and modulation of EEG activity, resulting in an increase of the spatial resolution of the EEG technique (Onton & Makeig, 2006).

Given *n* observed random variables *x*_1_, . . ., *x_n_*, which are a linear combination of *m* random source variables *s*_1_, . . ., *s_m_* as *x_i_* = *a_i_*_1_*s*_1_ + *a_i_*_2_*s*_2_ + *. . .* + *a_im_s_m_* for all *i*, where the *a_ij_* are real coefficients with *i* = 1, . . ., *n* and *j* = 1, . . ., *m*; the basic ICA model can be written as *x* = *As*, where *A* = [*a_ij_*] is a *n* × *m* mixing matrix, and each element *a_ij_* represents the mixing coefficient of the *j*-th source in the *i*-th observed mixture. Results of EEG data decomposition demonstrate that ICA, applied to sufficiently clean data, can separate around a dozen of maximally independent information sources whose scalp maps fit near-perfectly to dipolar projections of cortical EEG sources and non-brain signals like those related to eye movements, line noise, and muscle activity (Makeig, Debener, Onton, & Delorme, 2004).

The ICA approach assumes temporal independence among cortical sources. This assumption is sufficient to separate signals, independently of the physical distance among the brain sources (Onton & Makeig, 2006). The assumption of temporal independence is supported by neurophysiological evidence: the brain cortex is organized into compact regions of specialized function, where short connections are privileged over long connections (Delorme et al., 2012). Additional to cortical sources, an EEG recording also contains electrical activity related to eyes, muscles, defective electrodes, and line noise, but those sources have activity that is independent from the brain sources, agreeing with the ICA assumptions.

To compare the independent components (ICs) across different subjects two approaches are suggested: 1) clustering the ICs according to specific characteristics before the comparison (Onton & Makeig, 2006), making it difficult to establish that all the subjects under study have comparable ICs, or 2) the group ICA (gICA) analysis (Hyvärinen, 2013). One of the main gICA approaches is to concatenate the channels of different recordings, implying a common mixing matrix but different activations: [*x*_1_, *x*_2_, . . ., *x*_r_] = *A*[*s*_1_, *s*_2_, . . ., *s_r_*].

A first application of gICA in EEG was given by the work of Marco-Pallarés, Grau, and Ruffini (2005), where the brain sources related to the mismatch negativity ERP were found in two steps: a gICA decomposition and an inverse solution of the topography maps related to the ICs. In addition, Congedo, John, Ridder, Prichep, and Isenhart (2010) used an approach based in gICA to avoid the volume conduction effect over connectivity measures and stated that the approach is as useful in EEG as well as in fMRI. The same author, in a subsequent work, discussed how the gICA approach gives replicable physiological components that could help in diagnosis and assessment of abnormal brain functioning (Congedo, John, Ridder, & Prichep, 2010). The concatenation of subject data for gICA analysis has been applied to the study of Mild Cognitive Impairment (Ochoa et al., 2015), Alzheimer’s disease (Zervakis, Michalopoulos, Iordanidou, & Sakkalis, 2011) and Attention-Deficit Hyperactivity Disorder (Ponomarev, Mueller, Candrian, Grin-Yatsenko, & Kropotov, 2014).

The authors who prefer the clustering approach over the gICA approach emphasize the role of individual anatomical variability over the decomposition, opting to design methodologies that refine the clustering criterion (Bigdely-Shamlo, Mullen, Kreutz-Delgado, & Makeig, 2013). Some authors, alternatively, have introduced the Magnetic Resonance Image of the subjects (Tsai, Jung, Chien, Savostyanov, & Makeig, 2014), implying the loss of one of the principal advantages of EEG: its unbeatable lower cost compared to Magnetic Resonance Imaging techniques. The usage of gICA is encouraged by the work of Lio and Boulinguez (2018), who found that this approach is insensitive to inter-individual differences of neuroanatomy, and that the obtained spatial filters are optimized for analysis at the population level.

Considering the potential application of the gICA approach in searching for spatial filters that can be shared among subjects, various methodological approaches from individual ICA can be adapted to study the reliability of components identified using concatenated data. Approaches like ICASSO (Groppe, Makeig, & Kutas, 2009) have been used to evaluate the reliability of gICA decomposition using an ICA algorithm in the frequency domain (Labounek et al., 2018). Authors report fourteen stable components present across different paradigms (Labounek et al., 2018). More recently, the RELICA (Artoni, Menicucci, Delorme, Makeig, & Micera, 2014) was used on gICA components obtained from temporal and frequency algorithms, obtaining seven reliable components in control subjects using the frequency algorithm (Gholamipour & Ghassemi, 2021). The availability of different databases in the web, and the improvement in the approaches to combine them (Babayan et al., 2019; Bigdely-Shamlo et al., 2020), opens the possibility to study how different components could be reliably obtained from different databases.

In the current work, five databases from three different sites were analyzed using gICA to find common spatial filters among these databases. The gICA algorithm extracts spatial patterns common to comparable subjects in age using a common resting-state task, even when the site and protocol acquisition is different. The ICASSO and clustering methods were utilized to assess the reliability of the components, which led to the discovery of reproducible components across datasets. This opens up the possibility of establishing a baseline of spatial filters for studying neural activity in EEG recordings, based on gICA decomposition.

## 2 Materials and methods

### 2.1 Datasets

Five datasets were used to investigate the reproducibility of independent components:

#### 2.1.1 Alzheimer’s Biomarkers Dataset

This dataset comprises data from the project titled “Identification of preclinical biomarkers of Alzheimer’s disease through longitudinal monitoring of brain electrical activity in populations with a genetic risk” of the Neuropsychology and Behavior Group (GRUNECO, University of Antioquia) (García-Pretelt et al., 2022). Thirty healthy subjects (18 female, mean age of 34.8 ± 9.7, age range from 21 to 55) from the first longitudinal session of the project were used, specifically the eyes-closed 5- minute resting-state task. The dataset was acquired using the Neuroscan SynAmps2 amplifier with a 58-channel Electro-Cap following the “10-10” montage standard and referenced to a right earlobe electrode. Data was recorded at 1000 Hz and with a 0.1 Hz to 200 Hz bandpass. All participants gave written informed consent. The study protocol was approved by the bioethics committee of the Instituto de Investigaciones Médicas, Faculty of Medicine, Universidad de Antioquia, Colombia.

#### 2.1.2 Mind-Brain-Body Dataset

Corresponding to the Mind-Brain-Body Dataset of the Max Planck Institute (Babayan et al., 2019). A subset comprising 195 subjects (73 female, mean age of 39.3 ± 20.2, age range from 20 to 80) was used. Participants follow a resting-state task consisting of 16 alternating one-minute blocks of eyes-closed and eyes-open. The dataset was acquired using a “BrainAmp MR plus” amplifier in an electrically shielded and sound-attenuated EEG booth. Sixty-two channels were recorded using active Acti-CAP electrodes following the “10-10” montage standard and referenced to the FCz electrode. Data was recorded at a sampling frequency of 2500 Hz with a bandpass filter from 0.015 Hz to 1 kHz.

#### 2.1.3 Information Measures Dataset

This dataset consists of 22 subjects (11 female, 11 male, mean age = 21.1 ± 0.52 years, age range = 18-26) (L. Trujillo, 2019; L.T. Trujillo, Stanfield, & Vela, 2017). A resting state task was carried out with 4 minutes of eyes-open and 4 minutes of eyes-closed interleaved in 1-minute intervals. The dataset was acquired using the Bio Semi Active II amplifier with 72 active Ag/AgCl electrodes following the “10-5” montage standard and referenced to a common mode sense (CMS) electrode located between sites PO3 and POZ. Data was recorded at an initial sampling rate of 2,048 Hz (400-Hz bandwidth), and down sampled online to 256 Hz.

#### 2.1.4 Test-Retest Dataset

This dataset comprises data from Suárez-Revelo, Ochoa-G´ómez, and Tob´ón-Quintero (2018). It consists of 24 subjects (9 female, mean age of 31.4 ± 12.2, age range from 19 to 58). EEG signals were recorded at eyes-closed resting-state task during 5 minutes. A Neuroscan unit amplifier was used to record EEG signals. Data was recorded with a 0.05-200 Hz bandpass and sampled at 1000 Hz from 58 tin electrodes positioned according to the international “10-10” system with right earlobe reference.

#### 2.1.5 Precuneus Failures Dataset

This dataset consists of data acquired from the studies of Ochoa et al. (2017) and Duque-Grajales et al. (2014). From these works, 21 healthy subjects were collected (13 female, mean age of 30.37 years ± 5.7, age range from 19 to 41). A Neuroscan amplifier was used to record the EEG. Recordings were obtained while the subjects remained comfortably sat, resting with eyes closed for 5 minutes. EEG data were recorded from 64 electrodes (0.1–200Hz bandpass) with midline reference and sampling rate of 1000Hz. The electrodes were positioned according to the international “10–10” system.

### 2.2 Signal pre-processing

Each dataset was preprocessed with a pipeline inspired by the work of Suárez-Revelo et al. (2018), in order to improve test-retest reliability. The preprocessing pipeline used is explained in the following subsections and illustrated in Figure 1.

**Fig. 1.**
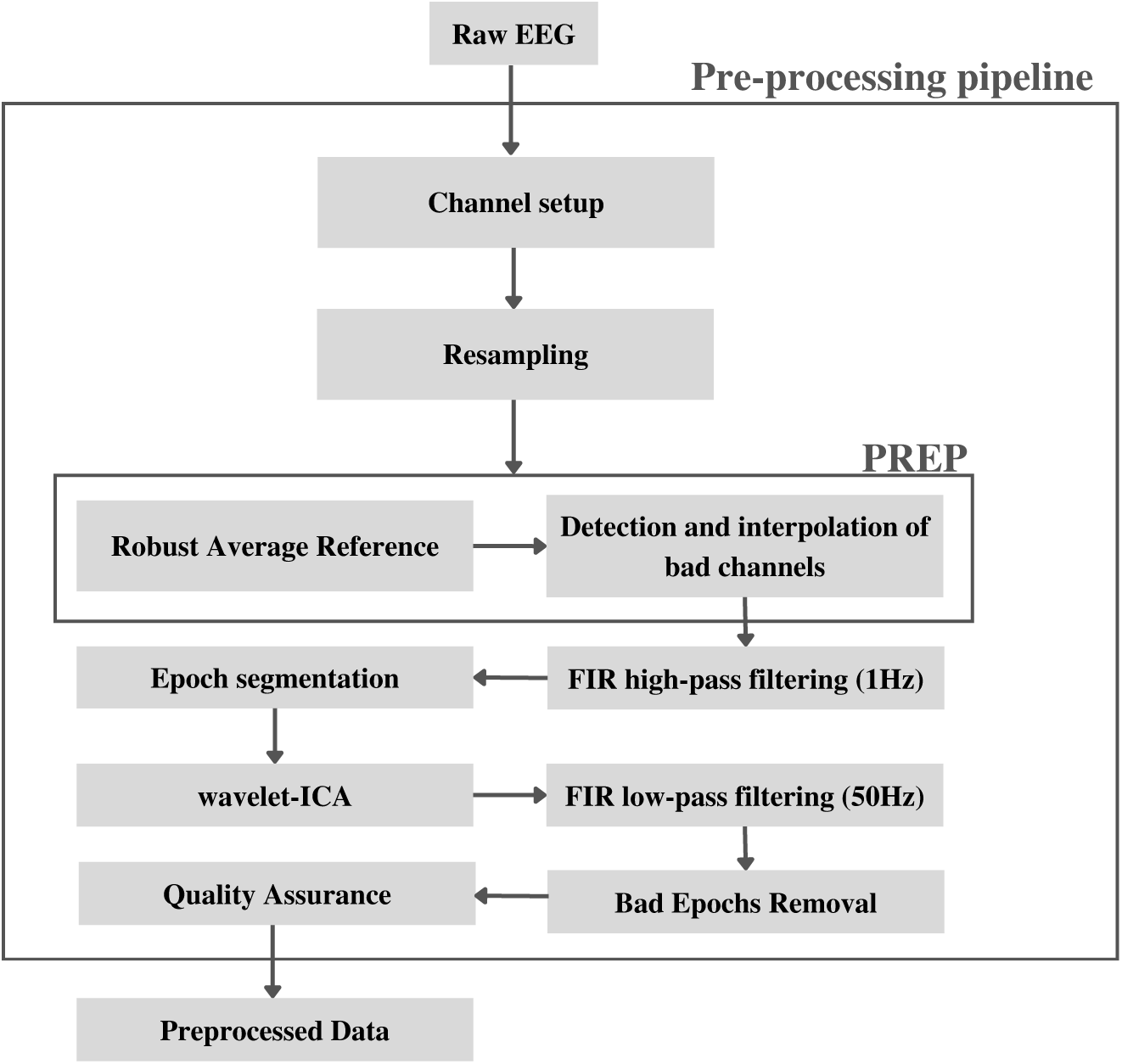
Diagram of the Pipeline for EEG Signal Pre-processing (Suárez-Revelo et al., 2018). The figure illustrates the step-by-step pipeline used for pre-processing EEG signals to enhance test-retest reliability.

#### 2.2.1 Channel Setup

For each dataset only the relevant EEG electrodes were selected. The specific electrodes used for each dataset can be found in Appendix B. The channel locations are set up by mapping the labels to the location defined on the “standard-10-5-cap385.elp” file of the DIPFIT plugin of EEGLAB (Delorme & Makeig, 2004, 2018).

#### 2.2.2 Resampling

To maintain a constant sampling frequency along the different datasets, each one was resampled to the minimum sampling frequency, which was 256 Hz.

#### 2.2.3 PREP pipeline

The PREP pipeline (Bigdely-Shamlo, Mullen, Kothe, Su, & Robbins, 2015) was applied on the continuous signal using the default parameters for the detection and interpolation of noisy channels, and for the computation of the robust average reference. For power line interference removal, the appropriate value (50 Hz or 60 Hz) was selected depending on the dataset. From this fundamental frequency, four harmonics are added (e.g., for 60 we get 120, 180, 240, 300). If any of these frequencies is larger than the Nyquist frequency, then the aliased frequency is considered if it is larger than the noise fundamental frequency. This last heuristic avoids the attenuation of the main EEG frequencies, commonly below the line noise.

#### 2.2.4 Epoch Segmentation

Resulting signals are segmented into epochs of 2 seconds to maintain stationarity (Gonen & Tcheslavski, 2012) and the non eyes-closed epochs are removed if there are any.

#### 2.2.5 wICA Correction

A detrending is done with a 1Hz high-pass filter implemented through the eegfilt-new function of EEGLAB v2021.0 (Delorme & Makeig, 2004) with default settings. Afterwards a decomposition was found using the FourierICA algorithm (Hyvärinen, Ramkumar, Parkkonen, & Hari, 2010) computed from 4Hz to 30Hz and with 30 PCA components. Subsequently, the Wavelet ICA (wICA) approach proposed by (Castellanos & Makarov, 2006) was applied to the FourierICA components found in order to correct artifacts.

#### 2.2.6 Epoch Rejection and Low Pass Filtering

Signals are then low passed to 50 Hz using the same eegfiltnew function to get the signal between 1Hz and 50Hz. Bad epochs are rejected using the pop autorej function of EEGLAB. This function first removes epochs with fluctuations above 1000uV and then iteratively removes epochs with join probability outside 5 standard deviations from the mean of the signal amplitude distribution (Onton & Delorme, 2008).

#### 2.2.7 Quality Assurance

Any EEG recording that does not follow any of the following requirements is rejected from the study:

- Have at least 90 clean epochs (3 minutes of data).
- Have at most 15% of interpolated channels in the prep pipeline.
- Have at most 5% of noisy channels in the prep pipeline.

### 2.3 Group Independent Component Analysis

The Group Independent Component Analysis explored in this study is based on the temporal concatenation of the EEG time-courses of N subjects. This assumes a similar mixing process among the subjects. Huster, Plis, and Calhoun (2015) recommended to use multi-level ICA when variability in the source topographies between subjects is to be expected. Nonetheless, here we are specifically interested in an averaged source topography along a group of subjects. Because of this, the approaches that return a single topography for each subject are not explored. An aggregation procedure could be done by clustering the individual topographies as suggested by Bigdely-Shamlo, Mullen, et al. (2013).

Independent Component Analysis is usually done on the time-domain. Nevertheless, a recent study by (Gholamipour & Ghassemi, 2021) shows that doing the analysis on the frequency-domain improves the reliability of the independent components found. Following this result, our study applies FourierICA (Hyvärinen et al., 2010). Importantly, one of the main parameters for ICA usage is the order of decomposition; here we explore a range of orders to select an optimal one as explained in section 2.4. Prior to the FourierICA, a dimensionality reduction procedure through PCA is applied using the chosen order. Another issue with ICA is that it may give different results across runs. This happens because of the different initialization values, the many local minima the objective function may have, and the iterative nature of the procedure. Consequently, the reliability of a component as intuitively defined by “does this component consistently appear across different runs of the algorithm?”, is not known from a single execution of the ICA. To account for this, the ICASSO methodology (Himberg & Hyvärinen, 2003) is applied by running the ICA algorithm as many times as there are subjects involved in the concatenation (N). The previous was done using a leave-one-out cross-validation scheme, that is, in each run one subject was excluded from the concatenated data. The results are then clustered to examine what components consistently arise from the data. Each cluster will have a representative component topography which is its centrotype or centroid (Himberg & Hyvärinen, 2003). Figure 2 illustrates the gICA procedure.

**Fig. 2.**
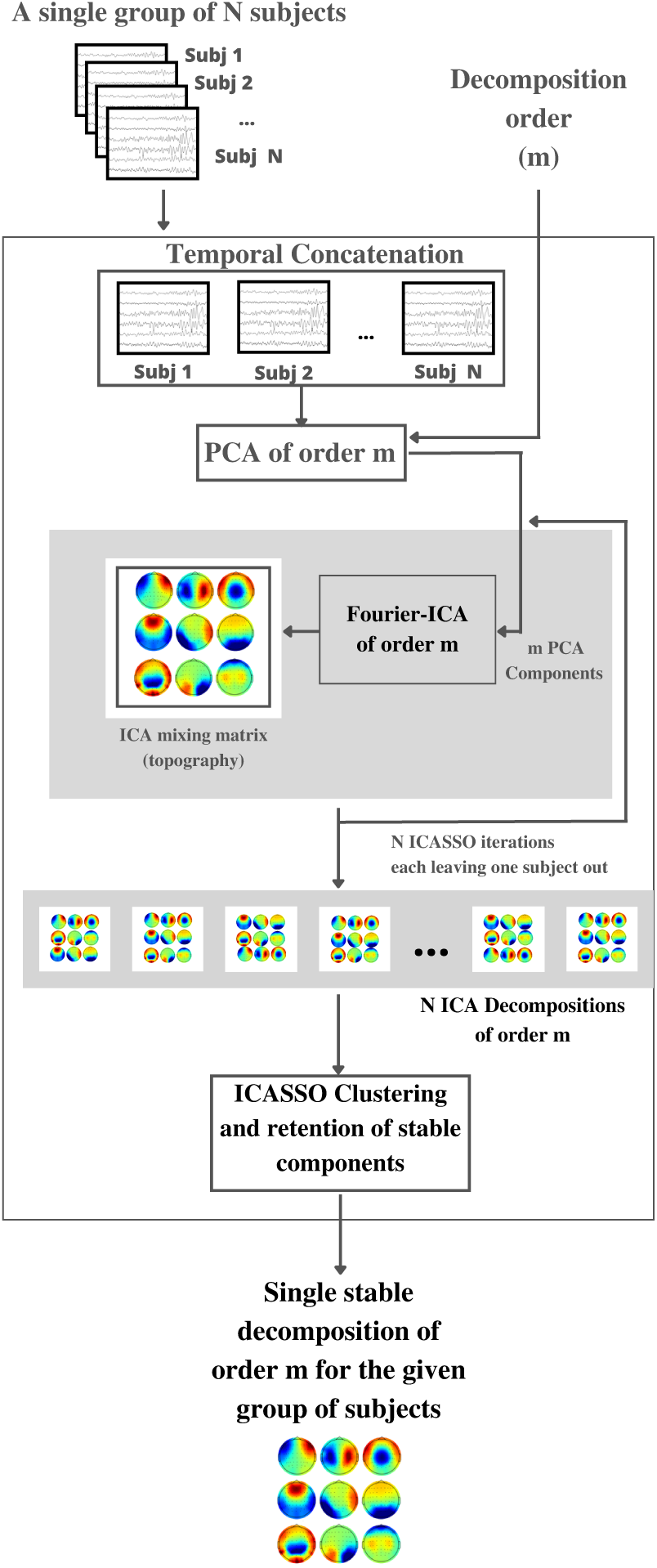
The gICA procedure has as input the EEG from N subjects, and a decomposition order m. The first step involves the temporal concatenation of the N subjects, followed by PCA of order m. Afterwards, the ICASSO methodology is implemented, that is, N iterations of Fourier-ICA are performed to evaluate reliability (noting that N is the number of subjects). In each of the iterations one subject is left out of the ICA concatenation input by slicing the concatenated array. This results in N ICA decompositions. Subsequently, the ICASSO clustering stage is conducted to obtain a single stable m-order decomposition for the given subjects.

### 2.4 Choosing the order of decomposition

One of the issues with ICA analysis is the estimation of the order of decomposition. Currently some analytical methods exist as those listed by Li, Adali, and Calhoun (2007). Nonetheless, a common finding among researchers (Miyakoshi, 2019) is that typically around 10 to 20 “good” independent components can be found using EEG data. Following this observation, here a more pragmatic approach is taken. A list of possible orders is generated from the range of 10 to 20 (inclusively), and three high order decompositions (25, 30 and 50) are added.

To select a suitable order of decomposition, an exploratory procedure was carried out. In essence, for a set of orders two kinds of metrics were calculated and then, from the curves of these metrics against the order, a single order was chosen for subsequent analyses. The metrics calculated are explained as follows:

#### 2.4.1 Reliability

In this first analysis, the main idea was to assess how reliable the gICA-inferred components were as a function of the order. Here, reliability is understood in the sense of the ICASSO methodology, that is, a component’s cluster is reliable if every run of the algorithm contributed to the cluster. Correspondingly, a cluster of components will lose reliability anytime a run of the gICA doesn’t contribute to its cluster. Nonetheless, we were not interested in the reliability of a single component but in the overall reliability of the components of a decomposition.

The following metrics were used as benchmarks of “reliability” for a single decomposition (formed by *N* clusters of components, where *N* is a given order):

Number of Reliable Clusters (NRC)

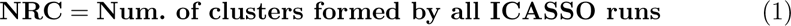

The higher the *NRC* the better, being the ceiling the total number of clusters which is equal to the order of decomposition.

- Reliability Ratio (RR)

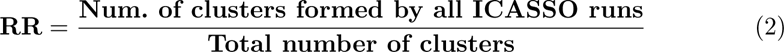

Notice that the “Reliability Ratio” will aggregate only the clusters formed by the same number of components as there were runs in the ICASSO.

This means that each run contributed one component to the cluster. Notice that the total number of clusters depends on the order, so the RR curve is NOT merely the *NRC* divided by a constant across the orders. Ideally, one would want this reliability ratio to be 1, which would mean every cluster accumulated a component in all the runs of the gICA algorithm in the ICASSO. The intuition we had regarding the relationship between order and reliability is that at a certain point they started to have an inverse relationship.

#### 2.4.2 Neural Likeness

Additionally, there is interest in searching for decompositions with a significant number of neural components. Classifying a component as neural is not a trivial matter; nevertheless, here we follow the ICLabel (Pion-Tonachini, Kreutz-Delgado, & Makeig, 2019) labelling method which estimates the probabilities of membership of a component to each one of 7 classes: Brain, Muscle, Eye, Heart, Line Noise, Channel Noise, and Other. ICLabel returns for each component a vector with 7 positions with each of the corresponding probabilities. A component will be classified as neural if the brain class is the one with the greatest probability. ICLabel does the classification from both the topography and the source signal demixed from the EEG. Although the temporally concatenated gICA infers the same topography for all subjects, the time-courses of the sources are different for each one of them; thus, a way to aggregate the results of ICLabel along subjects becomes necessary. Here the approach taken for this problem is to give the classifier the within-group subject-concatenated EEG signal as the time-course input and the given cluster centroid as topography (equivalent to the mean topography). Another strategy would be to do the labelling separately for each subject, then aggregate the results by summing the probabilities of each class along all the subjects, and finally find the class that maximized the sum. This was not explored for simplicity.

The following metrics are used as benchmarks of “neuralness” for a single decomposition (formed by *N* clusters of components, where *N* is a given order):

- Number of Neural Components (NNC)

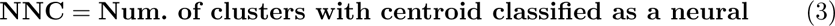

Naturally, one would want to maximize the *NNC*. Nonetheless, obtaining decompositions with only neural components is not expected since artifact related components arise. The intuition is that at a certain point the number of neural components stops increasing. This as the number of EEG-observable neural components has a limit (Miyakoshi, 2019).

- Mean Neural Probability (MNP)

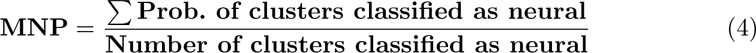

Notice that the NNC metric does not consider the probability that led to the cluster to be classified as neural. It is better to have less neural components with higher probability than the contrary. The “Mean Neural Probability” will thus consider the level of certainty. Ideally it should be as close to 1 as possible.

### 2.5 Group Formation

Now that the main metrics have been explained, we proceed to describe their application to the datasets. Noting that these metrics characterize a single decomposition (which is defined by the data used and the order), we segmented each dataset in groups of 15 subjects that shared a common age range (to avoid age-related effects) as illustrated in Table 1. The number of subjects was fixed to simplify the analysis since the quantity of data would increase the number of comparisons to be carried out in the order selection process. The only particularities for the group segmentation occurred in the “Test-Retest” dataset, which only has 9 participants in the “Adult” range (the other 15 are all Young Adults), so the group was completed with 6 subjects of the Adult range from the “Precuneus Failures” dataset. Lastly, the 15 participants left in the “Precuneus Failures” dataset were mostly from the “Young” age range, but three of these were slightly above (31, 32 and 33 years respectively). Appendix A details the resulting groups.

**Table 1.** Age Ranges for Group Formation.

| Label | Age Range |
| --- | --- |
| Young Adult | 18 - 30 |
| Adult | 31 - 59 |
| Old Adult | $\geq 60$ |

Following this procedure, 19 groups of 15 subjects were obtained (285 subjects in total). For all these groups a gICA decomposition was computed for each of the orders in the range mentioned before. Finally, the metrics described were calculated and, for each one, a distribution was made (from the results of the 19 groups). With the distributions, a single order is chosen to perform a clustering analysis.

### 2.6 Component Clustering

To assess the reproducibility of the components found through the groups, a corrmap-based clustering was carried out (Viola et al., 2009). In essence, the components are clustered based on their correlation to a template decomposition (see Appendix C). For a given gICA decomposition, each of its components is correlated to all the components of the templates, forming a 2D correlation matrix. Afterwards and through an iterative algorithm, the item of the matrix with the highest correlation is assigned to the corresponding component of the template. Then the procedure is repeated ignoring the previously assigned item and the component of the template it was assigned to. The procedure ends when all components of the template are filled. To further improve the quality of the clusters, a correlation threshold mask is applied to each of them. That is, components inside the cluster that do not meet a minimum correlation value (placed at 0.8) are removed from the cluster.

### 2.7 Cluster Characterization

After the clustering procedure, source localization is performed through dipole fitting of each component in each cluster. To this purpose, we used EEGLAB’s DIPFIT2 plugin using a template Boundary Element Model (Oostenveld & Oostendorp, 2002) extracted from the “Colin 27” canonical brain of the Montreal Neurological Institute (Holmes et al., 1998). For each cluster the mean dipole is obtained by averaging its Talairach coordinates (Talairach & Szikla, 1980). Afterwards, using the mean dipole coordinates, a Brodmann Area is associated to each cluster using the “Talairach Client” software (Brodmann, 1909; Lancaster et al., 1997, 2000). Similarly, for comparison purposes, this association was also obtained with the MNI2Tal tool from the Bio Image Suite (Lacadie, Fulbright, Arora, Constable, & Papademetris, 2008; Papademetris et al., 2006). Source activation time courses were extracted from the fully concatenated EEG signal (the time-wise concatenation of the EEG signals of all subjects in all groups); this required a reduction of channels to the montage intersection of all the datasets (which is available on Appendix B). The source activations were computed by building an ICA weight matrix from the average scalp maps of the clusters, and by performing the correspondent product with the concatenated EEG signal. Following the montage intersection, it was necessary to do a reordering/elimination of the channels in the ICA weights of the individual ICA decompositions of each group.

In addition, the clusters were characterized in the frequency domain by obtaining the mean power spectral density from the fully concatenated EEG signal mentioned earlier. This was done using the multitaper method implemented in MNE-python (Gramfort et al., 2013; Thomson, 1982). Furthermore, as was done during the order selection stage, each cluster centroid is classified into the 7 ICLabel classes using both the mean scalp maps of the clusters and the fully concatenated EEG signal as input. Finally, to measure the “reproducibility” of each cluster, two metrics were calculated:

- MTC (Mean Template Correlation): As explained before, the inclusion of a scalp map in a cluster depends on the correlation of that scalp map to a particular component of the selected template. This correlation must be above a threshold to be included. With this in mind, we define the mean template correlation of the cluster, that is, the average correlation value which supported the inclusion of the components to the corresponding cluster. As the cluster was formed with correlations above 0.8, this number serves as a baseline for the interpretation of this metric.
- APR (Appearance Ratio): The ratio of the number of groups that contributed to the cluster to the total number of groups studied. This can be interpreted as a ratio of appearance, or reproducibility.

To summarize the methodology, the diagram of Figure 3 serves as an overview of the whole procedure.

**Fig. 3.**
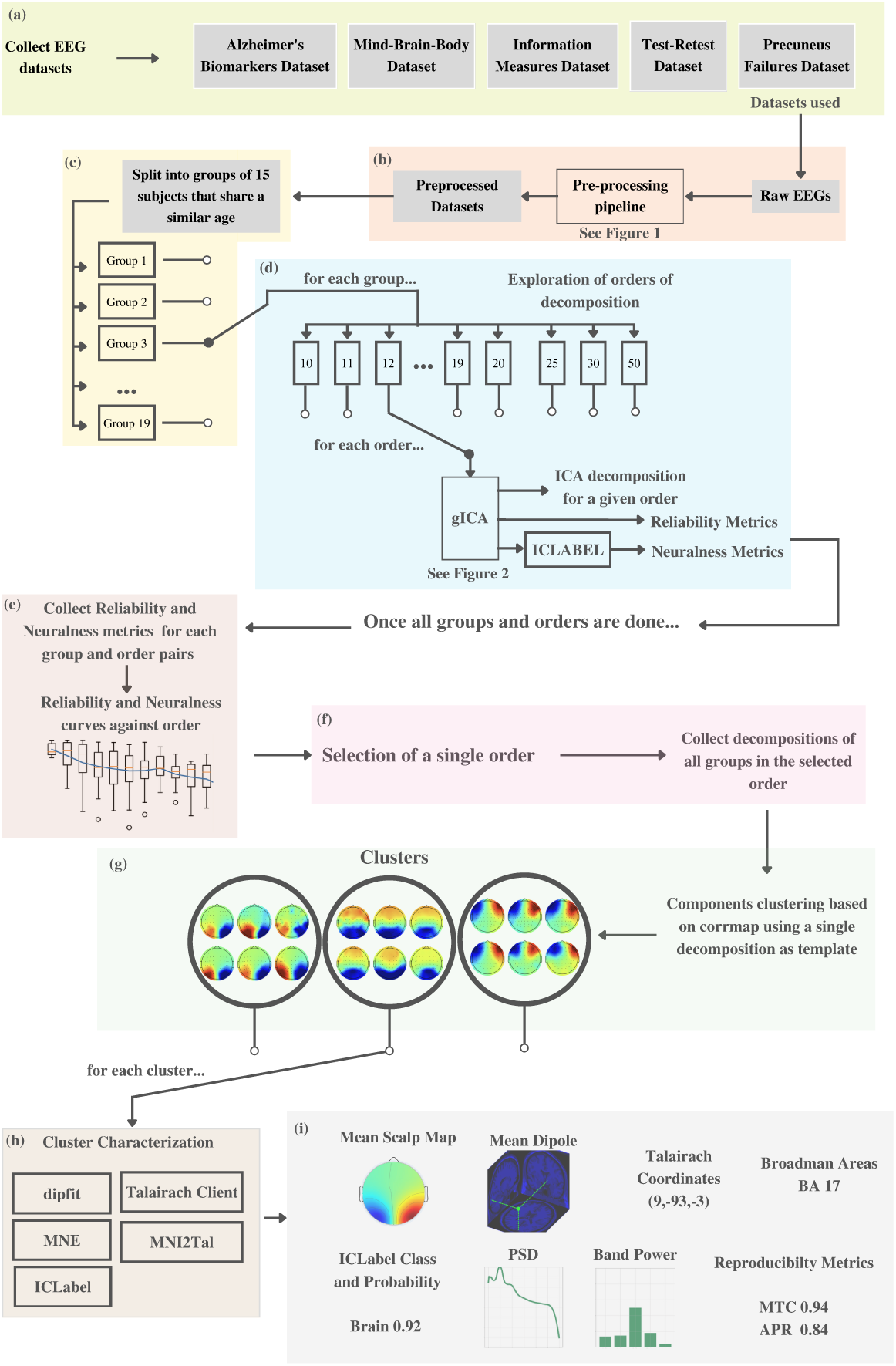
The methodology involves (a) collecting five datasets (from 3 different sites). Each EEG record from the dataset is (b) preprocessed. Subsequently, (c) groups of 15 subjects are formed maintaining a similar age within each group. (d) Each group will go through multiple gICAS [29] exploring different orders of decomposition, and each order is characterized by a set of reliability and neuralness metrics. After all the groups and orders are processed, (e) the information is collected and plotted in curves of the mentioned metrics against the order of decomposition. As a single order of decomposition is associated to multiple groups, boxplots are made to evaluate the generated distributions. From the visual inspection of these plots, (f) a single order of decomposition is chosen, prioritizing reproducibility, neuralness and a low variance across the groups. The data from the chosen order, representing multiple gICA decompositions (one by group), is collected to (g) perform a clustering of the components. The clusters are (h) characterized using different tools. Finally, the results include for each cluster (i) the mean scalp map, the mean dipole, its corresponding Talairach coordinates and Brodman areas, the ICLabel class and probability, the power spectral density, the band power, and the reproducibility metrics.

## 3 Results

### 3.1 Choosing the Order of Decomposition

Figure 4 shows the metrics described in section 2.4 plotted against varying orders of decomposition. From the point of view of “reliability” and “neuralness” the following observations can be made.

- The number of reliable components tends to be around ten until reaching higher orders (*>*30), where a higher number of reliable components is found as the order increases (Figure 4A).
- The percentage of reliable clusters diminishes as the order increases, demotivating the selection of higher orders, despite the previous observation (Figure 4C).
- The number of neural components tends to be around the dozens, even though some high numbers of neural components can be found in higher order (e.g. 50) decompositions (Figure 4B).
- Certainty of the neural components found diminishes as the order increases, and as before, this demotivates the use of higher order decompositions (Figure 4D).

**Fig. 4.**
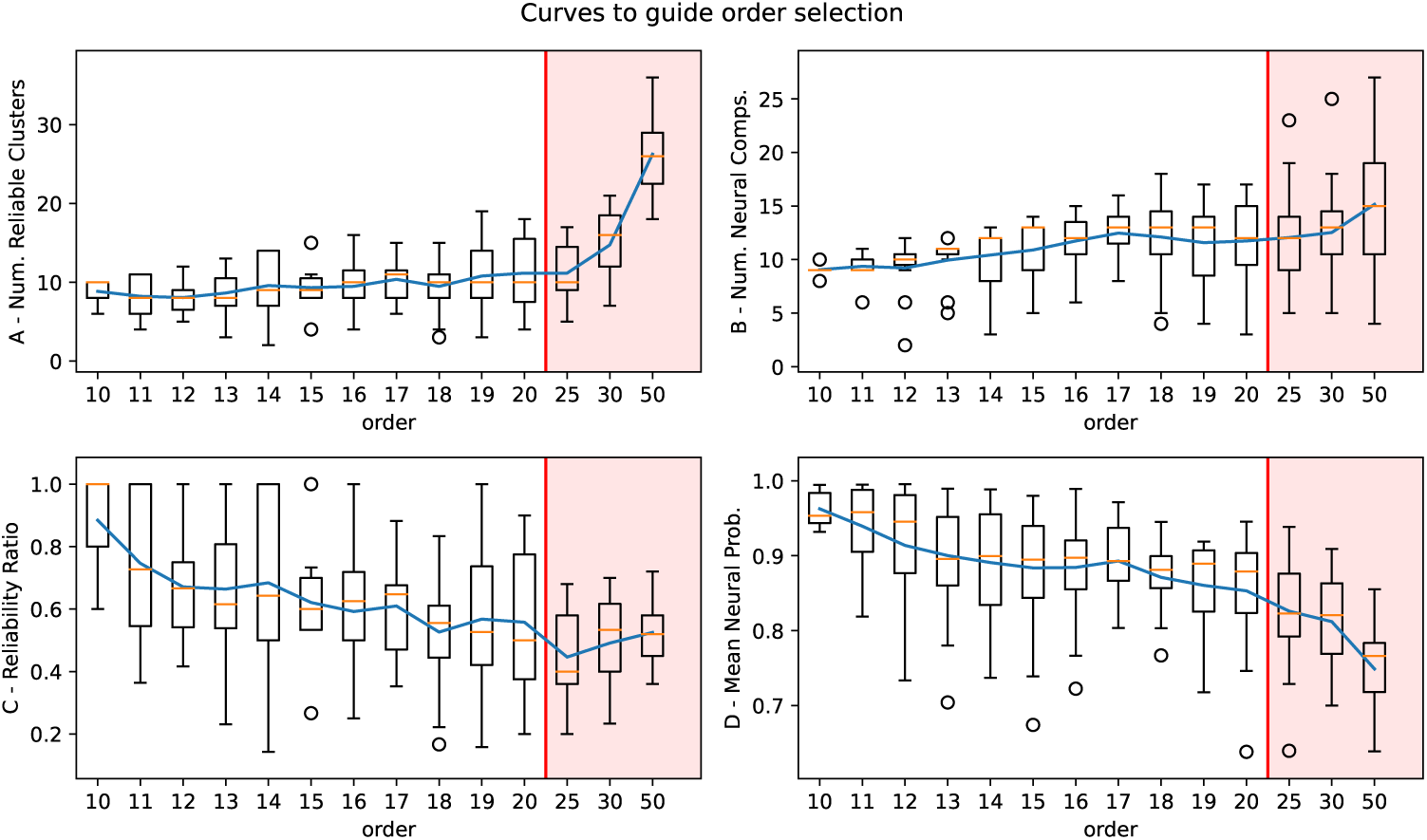
Curves of the metrics described in section 2.4 against the order of decomposition. For each order the distribution of the metric along the 19 gICA groups is plotted, and the blue curve is drawn through the mean of these distributions. The median is shown as the yellow line in each boxplot. The mildly red background is included as a warning of the curve not following a linear visualization, as the horizontal axis in the red section does not increase the order linearly.

Given the high dispersion of some of the distributions showcased in Figure 4, jointly with the low sample size (19 groups as points), the order chosen for the subsequent analyses was ten. This rationale is detailed as follows: First, the chosen order maintains the low dispersion along all metrics which assures that most of the individual samples keep the relatively good performance the selected order has both in terms of “neuralness” and “reliability”. And second, the previous proves to be important because if a higher but more dispersed order was chosen, later analyses would be greatly affected by the outlier points as the sample size is low. For example, order 11 showcases high dispersion in the Reliability Ratio, achieving bad performance (RR *<* 0.5).

### 3.2 Component Clustering

We aimed to investigate the reproducibility and characterization of ten components (C1-C10), each representing the centroid of their corresponding corrmap-derived clus-ter across the 19 groups with reliable decompositions obtained through the ICASSO methodology. The components identified from EEG data (which as mentioned before, are the centroids of the clusters), can be observed in Figure 5 sorted by APR (which is related to the reproducibility of the component). This figure shows topographic maps (Mean Cluster Scalp Map) which display the spatial distribution of each component across the scalp. Additionally, the individual (rather than the centroid) components inside the clusters are shown on Appendix D. Moreover, Figure 5 shows more information such as the 3D representation of the equivalent dipole of the cluster (Mean Dipole). Similarly, the Talairach coordinates (Mean Dipole Talairach Coords.) allows us to relate these components to specific Broadman areas (BAs) using the Talairach a Client and Brainsuite methods (denoted T and B, on Figure 5 respectively). Power spectral density (PSD) graphs showcase the frequency distribution of power associated with each component. Bar graphs depict the spectral power values within different frequency bands. To determine the component classes, we utilized ICLabel in Python, which provided probabilities of component classification (ICLabel). Additionally, the mean template correlation (MTC) values highlight the average correlation supporting component inclusion in their respective clusters, serving as a way to measure the internal consistency of the corresponding clusters. Lastly, the Appearance Ratio (APR) reveals the ratio between the number of contributing groups and the total number of groups studied. It is possible to cluster the components with the individual runs of the ICASSO rather than the resulting ICASSO decomposition, this is explored in Appendix E, with only minor changes to the main clusters.

**Fig. 5.**
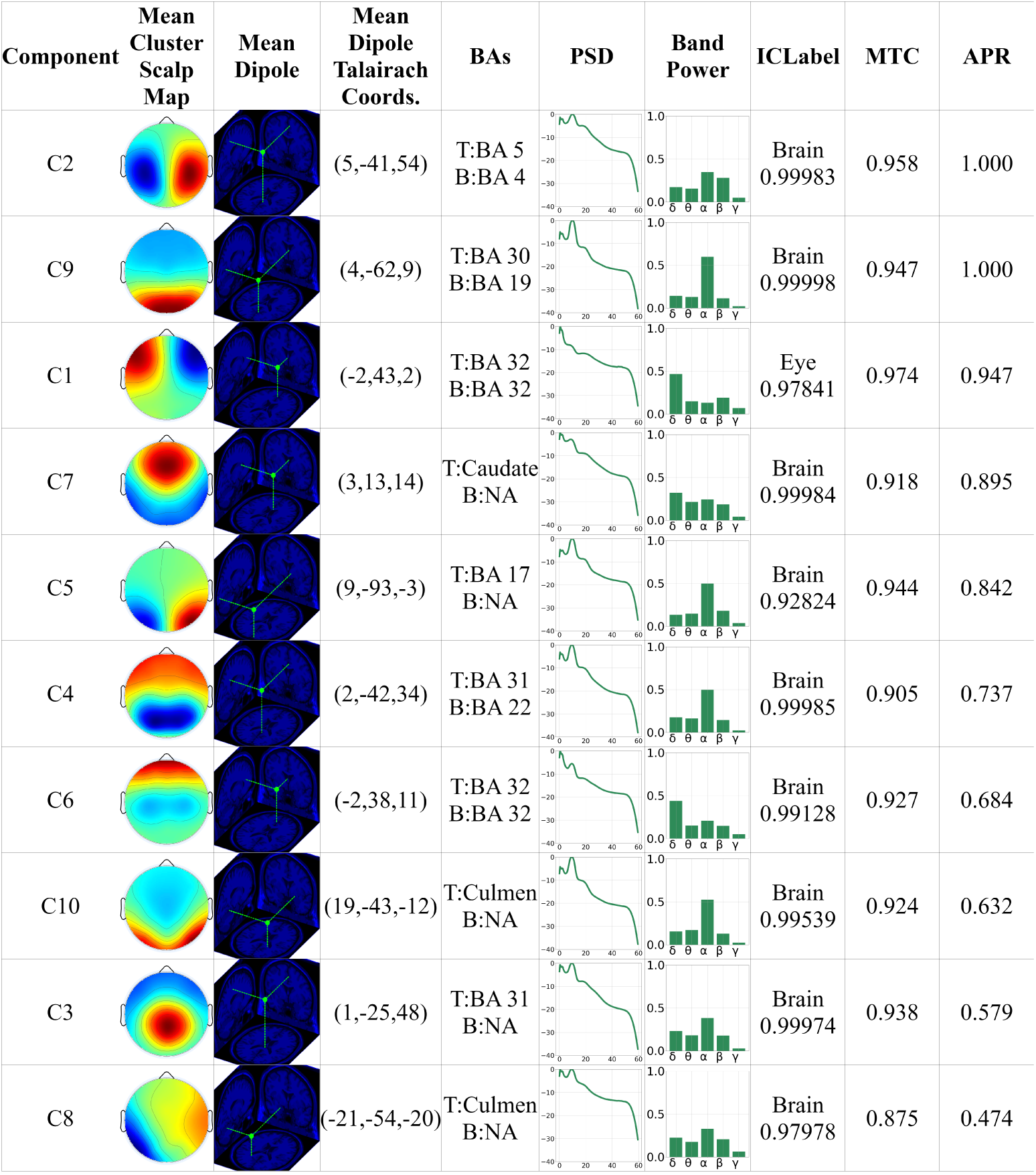
Localization, spectral, class and reproducibility characterization of EEG Components (C1- C10). Topographic maps (Mean Cluster Scalp Map) display their spatial distribution across the scalp. 3D representations (Mean Dipole) reveal estimated dipole locations and orientations. Talairach coordinates (Mean Dipole Talairach Coords) relate components to specific Broadman areas (BAs) using the Talairach Client and Brainsuite tools (T, B respectively). NA values indicate that the tool didn’t find a matching Brodman Area. Power spectral density (PSD) graphs showcase the frequency distribution of power associated with each component. Bar graphs depict spectral power values within different frequency bands. Component classification was done using ICLabel in Matlab, providing the associated class probabilities (ICLabel). Mean template correlation (MTC) values highlight average correlation of the components to the template supporting component inclusion in their respective clusters, showcasing how consistent are the clusters. Finally, the Appearance Ratio (APR) reveals the relationship between contributing components and the total groups studied, serving as a figure of merit for reproducibility. The table is sorted by this last column, having the most reproducible components at the top of the table.

The components shown in Figure 5 are described as follows:

#### 3.2.1 C1 Horizontal Eye Movement

Component C1 is located in the frontal lobe, specifically in the dorsal anterior cingulate cortex (dACC), Brodmann’s Area 32. However, it does not correspond to a neural source, and its appearance showed high reproducibility in some groups (APR=0.94) with a high MTC (over 95%). Its topography naturally resembles a horizontal EOG-related component, which is commonly found in the literature (Bigdely-Shamlo, Kreutz-Delgado, Kothe, & Makeig, 2013; Delorme et al., 2012; Hsu, Mullen, Jung, & Cauwenberghs, 2015; Radüntz, Scouten, Hochmuth, & Meffert, 2015; Wang, Chang, Chuang, & Liu, 2019; Wang, Ke, Chuang, & Liu, 2020).

#### 3.2.2 C2 Bilateral Motor

C2 is located in the Primary Motor Cortex and the Somatosensory Association Cortex (Brodmann’s Areas 4 and 5). It corresponds to a neural source, with high reproducibility across all groups (APR=1) and high consistent topographical distribution (MTC over 95%). In the frequency domain, C2 exhibits an alpha peak and substantial beta activity, consistent with motor oscillations in the literature (Baker, 2007; Barone & Rossiter, 2021; Delorme et al., 2012; Tsai et al., 2006). Moreover, this component can be found either combined or split into its left and right parts (Artoni et al., 2014; Delorme et al., 2012; Hsu et al., 2015; Lin, Shaw, young Young, Lin, & Jung, 2012; Radüntz et al., 2015; Sita & Nair, 2013; Tsai et al., 2006; Wang et al., 2019, 2020).

#### 3.2.3 C3 Central-Parietal

C3 is located in the dorsal posterior cingulate cortex (dPCC), Brodmann’s Area 31; part of the brain’s posterior cingulate gyrus in the medial cortex. It appears to be a neural source based on its location, 1/f spectrum, alpha peak, and ICLabel’s “brain” label. However, the component indicates low reproducibility (APR=0.58) and a high MTC of 94% suggests limited similarity between the template and the scalp map. Similar components related to brain information have been previously reported (Delorme et al., 2012; García-Pretelt et al., 2022; Gholamipour & Ghassemi, 2021; Ochoa et al., 2017). In the frequency domain, C3 shows an alpha peak and considerable beta activity. Some literature reports similar components as central, or fronto-central (Beldzik, Domagalik, Gawlowska, Marek, & Mojsa-Kaja, 2017; Lin et al., 2012; Töllner et al., 2017) or centro-parietal (Chen, Ros, & Gruzelier, 2013; Tsai et al., 2006).

#### 3.2.4 C4 Parietal

C4 is located around Brodmann’s Areas 22 and 31, known as the superior temporal gyrus and the dorsal posterior cingulate cortex (dPCC), respectively. Area 22 is involved in auditory processing and language comprehension, while Area 31 is associated with memory, spatial navigation, self-referential thinking, and emotional processing. The characteristics shown in Figure 5 suggest that C4 corresponds to a neural source. However, its appearance indicates medium reproducibility (APR=0.73), and a high MTC of 90% suggests limited consistency in its topographic distribution across groups. In the frequency domain, C4 exhibits an alpha peak and considerable beta activity, consistent with motor oscillations. Similar components can be found in the literature described as parietal components (Beldzik et al., 2017; Chen et al., 2013; Lai et al., 2012; Lin et al., 2012; Wang et al., 2019, 2020). Others as a posterior alpha component (Artoni et al., 2014; Delorme et al., 2012; Tsai et al., 2006), or without label (Bian, Wang, Cao, & Zhang, 2006; Gholamipour & Ghassemi, 2021; Radüntz et al., 2015).

#### 3.2.5 C5 Bilateral Occipital

C5 is located around Brodmann’s Area 17, the primary visual cortex (V1), in the occipital lobe. V1 is a critical region for visual perception, being the first cortical area in the visual processing pathway. The characteristics of C5, including its location, 1/f spectrum, alpha peak, and ICLabel’s “brain” label, suggest that it corresponds to a neural source. However, the component indicates medium reproducibility (APR=0.84), and a high MTC of 94% suggests limited similarity between the template and the scalp map. Similar components related to brain information are commonly found in literature characterized as occipital activity (Chen et al., 2013; Lai et al., 2012; Wang et al., 2019, 2020).

#### 3.2.6 C6 Frontal

C6 is located around Brodmann’s Area 32, the dorsal anterior cingulate cortex (dACC), in the frontal lobe. It is part of the medial prefrontal cortex, specifically in the anterior cingulate gyrus. The characteristics of C6, such as its location, 1/f spectrum, alpha peak, and ICLabel’s ”brain” label, suggest it corresponds to a neural source. However, the component appeared with low reproducibility (APR=0.68), and a high MTC of 93% suggests limited similarity between template and the scalp map. Similar components related to blinking are commonly found in literature (Artoni et al., 2014; Bigdely-Shamlo, Kreutz-Delgado, et al., 2013; Delorme et al., 2012; Radüntz et al., 2015), whereas others do not explicitly mention this, effectively characterizing it as a frontal component (Castiglione, Wagner, Anderson, & Aron, 2019; Chen et al., 2013; Gholamipour & Ghassemi, 2021; Onton & Makeig, 2006; Sita & Nair, 2013; Wang et al., 2020).

#### 3.2.7 C7 Frontal Medial

C7 is a neural source located around the caudate nucleus, a subcortical structure in the basal ganglia. It plays a crucial role in motor control, reward processing, learning, and executive function. The component appeared with high reproducibility (APR=0.89), and a high MTC of 91% indicates similarity between brain data and the scalp map. The theta peak is more significant than the alpha peak, aligning with previous studies exploring theta rhythm in cortices (Delorme et al., 2012; Lai et al., 2012). Similar components are frequent in the literature Artoni et al. (2014); Castiglione et al. (2019); Chen et al. (2013); Delorme et al. (2012); Hsu et al. (2015); Lai et al. (2012); Lin et al. (2012); Onton and Makeig (2006); Tsai et al. (2006); Töllner et al. (2017); Wang et al. (2019).

#### 3.2.8 C8 Bilateral Temporal

C8 is located around the culmen, a part of the cerebellum, which is a separate structure from the cerebral cortex. The cerebellum is involved in motor coordination, balance, and posture. The location, 1/f spectrum, alpha peak, and ICLabel’s almost certain “brain” label suggest that C8 corresponds to a neural source. The component appeared in some of the groups, indicating a low reproducibility of (APR=0.47). Similarly, the medium MTC of 87% indicates that there is relatively little similarity or correspondence between the template and the scalp map. Similar components can be found in literature (Wang et al., 2019, 2020) split into its left and right parts, others associate the bilateral pattern with the electrocardiogram (Radüntz et al., 2015).

#### 3.2.9 C9 Occipital

C9 corresponds to a neural source located around Brodmann’s Areas 30 and 19. Area 30 is associated with emotional processing and mood regulation, while Area 19 is involved in higher-level visual processing and visual perception. The component showed high reproducibility (APR=1) across all groups, and the topographic distribution was consistent (MTC=95%) among them. The location, 1/f spectrum, alpha peak, and ICLabel’s “brain” label all support its neural nature. Similar components related are commonly found in literature (Chen et al., 2013; Lai et al., 2012; Lin et al., 2012; Sita & Nair, 2013; Wang et al., 2019, 2020).

#### 3.2.10 C10 Occipital-Temporal

C10 corresponds to a neural source located around the culmen, a part of the cerebellum responsible for motor coordination, balance, and posture. The component’s location, 1/f spectrum, alpha peak, and ICLabel’s ”brain” label all support its neural nature. It appeared with low reproducibility (APR=0.63), and a high MTC of 92% suggests limited consistency in its topographic distribution across groups. Similar components can be found in the literature, combined or separated into its left and right parts (Artoni et al., 2014; Chen et al., 2013; Onton & Makeig, 2006; Sita & Nair, 2013).

## 4 Discussion

The primary objective of this study was to evaluate the effectiveness of Group Independent Component Analysis (gICA) in decomposing EEG data for accurate identification of reproducible neural sources. Employing five distinct datasets with a total of 292 healthy subjects on the resting-state condition, the gICA methodology was applied individually to 19 groups of 15 subjects across different orders of decomposition. To continue our analysis, a single one of the orders had to be chosen given the computational limitations and the complexity of the results of subsequent stages. The process of estimating the correct number of sources or components within a dataset is a complex endeavor. It depends on factors such as data complexity and the nature of the neural sources. Mismatches between the chosen model and the actual sources can lead to suboptimal results, underscoring the need for precise model selection (Hyvärinen, 2013). We approached the problem by characterizing them in regards to reliability and neuralness, and from this it was found that lower orders exhibit better properties. This agrees with the observation of Miyakoshi (2019), thus higher decomposition orders don’t prove to be useful.

After choosing a low order of decomposition (10), the components found across the groups were clustered and characterized by different properties but specially by how common was to find them (APR metric), and how consistent was with the overall cluster (MTC metric). Overall, we identified ten components with varying levels of reproducibility, which represent spatial filters. This opens up the possibility of using the obtained decomposition as a prior for inverse solution methods (Lei, Wu, & Valdes-Sosa, 2015), as a constraint for further ICA decomposition (De Vos, De Lathauwer, & Van Huffel, 2011), or as a reference pattern for new approaches in designing spatial filters based on deep learning (Joo, Quan, Kim, Woo, et al., 2023). Moreover, their correlation to other spatiotemporal maps could be studied, such as those derived from microstates (Michel & Koenig, 2018). Of the ten components, around nine can be classified as neural. In particular, the bilateral motor (C2), frontal medial (C7), and occipital (C9) neuronal components were the most reproducible across the groups, appearing in more than 89% of them. Other components are moderately reproducible such as C4 and C5. Finally, some components appeared in less than 70% of the groups (C6,C10,C3,C8). Each of these components can be found on the literature in varying numbers as described in the results. Nevertheless, the comparison of components across the different literature involves difficult judgements. For example, bilateral components (such as C2) can be found split or combined into their left and right parts something that might related to the order of decomposition.

Additionally, verbally describing these components present inconsistencies among the literature. For example, C4 (which we describe as a parietal component), can be found as a posterior alpha component (Artoni et al., 2014; Delorme et al., 2012; Tsai et al., 2006). Others, have a similar parietal label (Beldzik et al., 2017; Chen et al., 2013; Lai et al., 2012; Lin et al., 2012; Wang et al., 2019, 2020), and naturally some don’t label the component at all (Bian et al., 2006; Gholamipour & Ghassemi, 2021; Radüntz et al., 2015). The labeling procedure is subjective, and it is difficult to assess the exact position of the component. Usually components with similar patterns with different relative positions with respect to the topography borders can be seen. Moreover, characteristics that would further identify the components, such as dipole locations and the frequency spectrum, are not necessarily reported along the topography.

Regarding the internal consistency of the topographies inside each cluster, evaluating the mean correlation of the components of the clusters to the template (MTC) didn’t prove to be useful, as the values are all high (over 80%). This comes naturally from the fact that the threshold for acceptance into a cluster was 0.8. Nevertheless, it is apparent that the mean MTC of the top-5 reproducible components (0.948, components sorted by APR) is greater than the mean of the bottom-5 ones (0.913), showcasing that indeed the most reproducible components tend to group into clusters with more consistent topographies. For example, C8 showcases both the minimum reproducibility (lowest APR) and the minimum consistency of the cluster (lowest MTC).

Apart from neural components, some noise sources can be highly reproducible and consistent, such as those reflecting horizontal eye activity (C1). This showcases the importance of addressing of common artifacts. Besides discussing the components, methodologically there still exists unexplored approaches such as the concatenation of subjects from different datasets, but this would prove to be of limited usage as usually a single dataset would have enough data to fill the available computational memory. In this sense, a pure cluster-based approach (without subject concatenation) such as the one proposed by Bigdely-Shamlo, Mullen, et al. (2013) may be more appropriate. Nevertheless, given that ICA decompositions tend to improve with data availability, the methodology proposed here, consisting of subject-concatenation into groups and a later clustering, could represent a good alternative when lengthy recordings are not available at the subject level.

It is crucial to acknowledge the study’s limitations. One of this is the variability in EEG source topographies, which can arise from variations in electrode placement among subjects or differences in brain anatomy. This phenomenon poses a particular challenge for algorithms that rely on the assumption of a common mixing matrix between the subjects (Hyvärinen, 2013). Similarly, high variability in the onset times of neural responses is frequently encountered in studies involving evoked responses. This variation can hamper algorithm performance, complicating data alignment and the extraction of consistent components(Hyvärinen, 2013). Moreover, although an order of decomposition of ten for all groups was methodologically grounded by selecting it according to the reliability and neuralness characteristics, it could be that different groups have different optimal orders of decomposition. In this work we constraint all groups to have the same order. Thus, there still challenges associated with order estimation. Finding the optimal order per group and then clustering might be an alternative, but would require a more refined selection of the template for the use of corrmap, or using another clustering approach.

Moreover, although a high number of subjects (292) were used in the current study, the sample size of the groups is low (19). Additional work is needed to collect more datasets to improve this sample size. Importantly, the site of origin is an important factor to control in subsequent works, as the present results might be affected by the overrepresentation of the largest dataset.

Overall, this study underscores the potential and challenges of using gICA for EEG data decomposition. While we identified several reproducible neural components, reflecting a broad spectrum of brain activities, our findings also bring to light the intricacies of EEG analysis, particularly in order selection and the interpretation of neural sources. Despite the limitations posed by sample size and dataset variability, our results provide valuable insights into reproducible neural sources appearing in resting-state conditions, enriching our understanding of brain function obtained using EEG. Moreover, the methodology provided can be particularly useful for researchers interested in leveraging the multi-subject nature of the gICA approach, offering a valuable framework for future EEG research. As the field advances, further studies with larger and more diverse datasets are essential for refining these techniques and deepening our understanding of neural dynamics.

## 5 Conclusions

This study aimed to evaluate the effectiveness of Group Independent Component Analysis (gICA) in identifying reliable neural components from resting-state EEG datasets across multiple sites. Our analysis of five databases covering 292 healthy subjects segmented into 19 groups revealed that lower order decompositions are more beneficial in the gICA methodology. By choosing an order of ten, nine neural components were identified, being the most reproducible the bilateral motor, frontal medial, and occipital neuronal components, appearing in more than 89% of the groups.

The employed methodology allowed for the identification of reproducible neural components which may be used as spatial filters between different datasets, indicating its potential for broad application in EEG research. However, the study also highlighted challenges in EEG data analysis, such as the variability of EEG source topographies, complexities in model order selection, and low group-wise sample size, underscoring the need for refined techniques and larger, more diverse datasets for future studies.

Overall, this research provides valuable insights into the reproducibility of neural sources in resting-state conditions and offers a robust framework for future EEG analyses. It emphasizes the potential and limitations of using gICA for EEG data decomposition and underlines the importance of further studies to advance our understanding of neural dynamics. Potential avenues for future work include exploring the use of spatial filters obtained through gICA for the extraction and analysis of brain activations, along with investigating the relation between these components and other spatiotemporal maps. These exploratory directions could significantly enrich our comprehension of brain activity patterns and enhance the methodologies in EEG analysis.

## 6 Statements and Declarations

## Funding

This work was supported by Ministerio de Ciencia, Tecnología e Innovación (MIN-CIENCIAS) through the project “Identificación de Biomarcadores Preclínicos en Enfermedad de Alzheimer a través de un Seguimiento Longitudinal de la Actividad Eléctrica Cerebral en Poblaciones con Riesgo Genético”, identified with the code 111577757635.

Similarly, the authors acknowledge the support provided by Comité para el Desarrollo de la Investigación - CODI Universidad de Antioquia, through the project “Cambios en los patrones del electroencefalograma cuantitativo (reactividad alfa, theta y su índice) en reposo y tareas de memoria, en el seguimiento longitudinal de pacientes con riesgo genético para Enfermedad de Alzheimer Temprano”, identified with the code 2017-16371.

## Competing interests

None of the authors declared any conflict of interest.

## Ethics approval

Not applicable

## Consent to participate

Not applicable

## Consent for publication

Not applicable

## Availability of data and materials

Data are available upon reasonable request.

## Code availability

Scripts are available upon reasonable request.

## Authors’ contributions

**A:** Conceptualization. **B:** Methodology. **C:** Software. **D:** Validation. **E:** Formal analysis. **F:** Investigation. **G:** Resources. **H:** Data Curation. **I:** Writing - Original Draft. **J:** Writing - Review Editing. **K:** Visualization. **L:** Supervision. **M:** Project administration. **N:** Funding acquisition.

John Fredy Ochoa-Gomez **A,B,C,I,J,L,M**

Yorguin-Jose Mantilla-Ramos **B,C,E,F,G,H,I,J,K**

Veronica Henao Isaza **D,I,J,K**

Carlos Andres Tobon **G,N**

Francisco Lopera **G,N**

David Aguillon **G,N**

Jazmin Ximena Suarez **B,G,I,J,L**

## Appendix A Table of Groups

| Group Number | Dataset Source | Age Range | Group Label |
| --- | --- | --- | --- |
| 1 | Alzheimer's Biomarkers Dataset | Adult | BALZ-A1 |
| 2 | Alzheimer's Biomarkers Dataset | Young | BALZ-Y1 |
| 3 | Precuneus Failures Dataset | Young<br>(3/15 adults) | PFCA-Y1 |
| 4 | Mind-Brain-Body Dataset | Adult | MBBD-A1 |
| 5 | Mind-Brain-Body Dataset | Old | MBBD-O1 |
| 6 | Mind-Brain-Body Dataset | Old | MBBD-O2 |
| 7 | Mind-Brain-Body Dataset | Old | MBBD-O3 |
| 8 | Mind-Brain-Body Dataset | Old | MBBD-O4 |
| 9 | Mind-Brain-Body Dataset | Young | MBBD-Y1 |
| 10 | Mind-Brain-Body Dataset | Young | MBBD-Y2 |
| 11 | Mind-Brain-Body Dataset | Young | MBBD-Y3 |
| 12 | Mind-Brain-Body Dataset | Young | MBBD-Y4 |
| 13 | Mind-Brain-Body Dataset | Young | MBBD-Y5 |
| 14 | Mind-Brain-Body Dataset | Young | MBBD-Y6 |
| 15 | Mind-Brain-Body Dataset | Young | MBBD-Y7 |
| 16 | Mind-Brain-Body Dataset | Young | MBBD-Y8 |
| 17 | Test-Retest Dataset | Young | TEST-Y1 |
| 18 | Test-Retest Dataset<br>(6/15 from Precuneus Failures Dataset) | Adult | TEST-A1 |
| 19 | Information Measures Dataset | Young | INME-Y1 |

## Appendix B Channel selection for each dataset

- Alzheimer’s Biomarkers Dataset - 58 channels: AF3, AF4, C1, C2, C3, C4, C5, C6, CP1, CP2, CP3, CP4, CP5, CP6, CPZ, CZ, F1, F2, F3, F4, F5, F6, F7, F8, FC1, FC2, FC3, FC4, FC5, FC6, FCZ, FP1, FP2, FPZ, FZ, O1, O2, OZ, P1, P2, P3, P4, P5, P6, P7, P8, PO3, PO4, PO5, PO6, PO7, PO8, POZ, PZ, T7, T8, TP7, TP8
- Mind-Brain-Body Dataset - 61 channels: AF3, AF4, AF7, AF8, AFZ, C1, C2, C3, C4, C5, C6, CP1, CP2, CP3, CP4, CP5, CP6, CPZ, CZ, F1, F2, F3, F4, F5, F6, F7, F8, FC1, FC2, FC3, FC4, FC5, FC6, FP1, FP2, FT7, FT8, FZ, O1, O2, OZ, P1, P2, P3, P4, P5, P6, P7, P8, PO10, PO3, PO4, PO7, PO8, PO9, POZ, PZ, T7, T8, TP7, TP8
- Information Measures Dataset - 64 channels: AF3, AF4, AF7, AF8, AFZ, C1, C2, C3, C4, C5, C6, CP1, CP2, CP3, CP4, CP5, CP6, CPZ, CZ, F1, F2, F3, F4, F5, F6, F7, F8, FC1, FC2, FC3, FC4, FC5, FC6, FCZ, FP1, FP2, FPZ, FT7, FT8, FZ, IZ, O1, O2, OZ, P1, P10, P2, P3, P4, P5, P6, P7, P8, P9, PO3, PO4, PO7, PO8, POZ, PZ, T7, T8, TP7, TP8
- Test-Retest Dataset - 58 channels: AF3, AF4, C1, C2, C3, C4, C5, C6, CP1, CP2, CP3, CP4, CP5, CP6, CPZ, CZ, F1, F2, F3, F4, F5, F6, F7, F8, FC1, FC2, FC3, FC4, FC5, FC6, FCZ, FP1, FP2, FPZ, FZ, O1, O2, OZ, P1, P2, P3, P4, P5, P6, P7, P8, PO3, PO4, PO5, PO6, PO7, PO8, POZ, PZ, T7, T8, TP7, TP8
- Precuneus Failures Dataset - 62 channels: AF3, AF4, C1, C2, C3, C4, C5, C6, I1, I2, CP1, CP2, CP3, CP4, CP5, CP6, CPZ, CZ, F1, F2, F3, F4, F5, F6, F7, F8, FC1, FC2, FC3, FC4, FC5, FC6, FCZ, FP1, FP2, FPZ, FT7, FT8, FZ, O1, O2, OZ, P1, P2, P3, P4, P5, P6, P7, P8, PO3, PO4, PO5, PO6, PO7, PO8, POZ, PZ, T7, T8, TP7, TP8
- Intersection - 54 channels: AF3, AF4, C1, C2, C3, C4, C5, C6, CP1, CP2, CP3, CP4, CP5, CP6, CPZ, CZ, F1, F2, F3, F4, F5, F6, F7, F8, FC1, FC2, FC3, FC4, FC5, FC6, FP1, FP2, FZ, O1, O2, OZ, P1, P2, P3, P4, P5, P6, P7, P8, PO3, PO4, PO7, PO8, POZ, PZ, T7, T8, TP7, TP8

## Appendix C Template used for clustering

One of the icasso decompositions of one of the 15-subject groups studied was used as a template for the corrmap clustering. The template was chosen by visual inspection, giving priority to the decompositions where each one of the components was clearly identifiable.

**Fig. C1.**
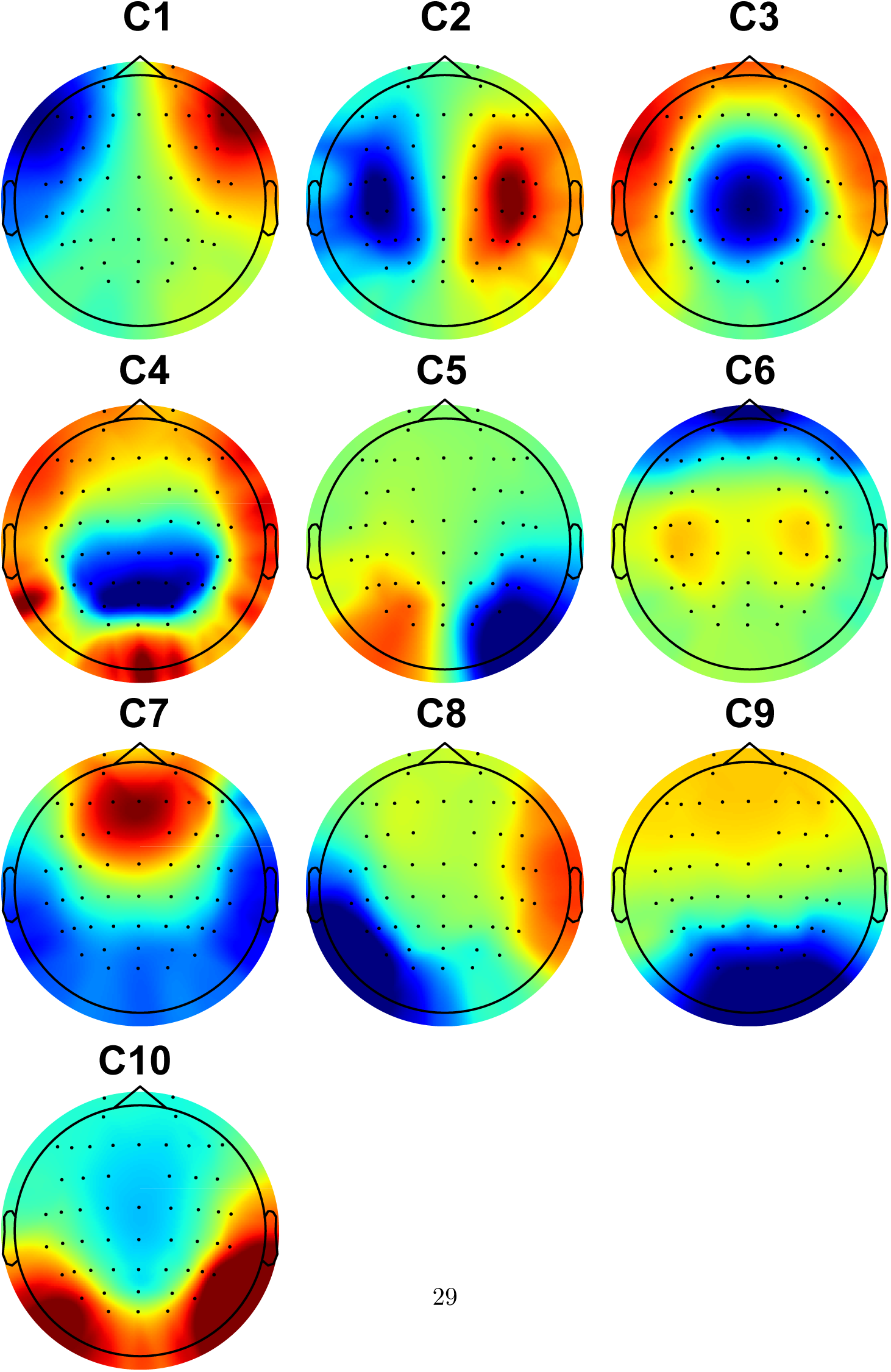
The template used for the corrmap clustering procedure.

## Appendix D Individual decompositions inside of each cluster

To give a complete account of the clustering results, the individual components that were grouped into each cluster are presented in the following figures. TC stands for the “template correlation” which is the correlation between the corresponding component of the template and the specific scalp map evaluated. APR is the appearance ratio and MTC is Mean Template Correlation, as defined in the main text. The first scalp-map corresponds to the Mean Scalp Map of the cluster. The second one is the one of the template.

### D.1 Component 1

See Figure D2.

**Fig. D2.**
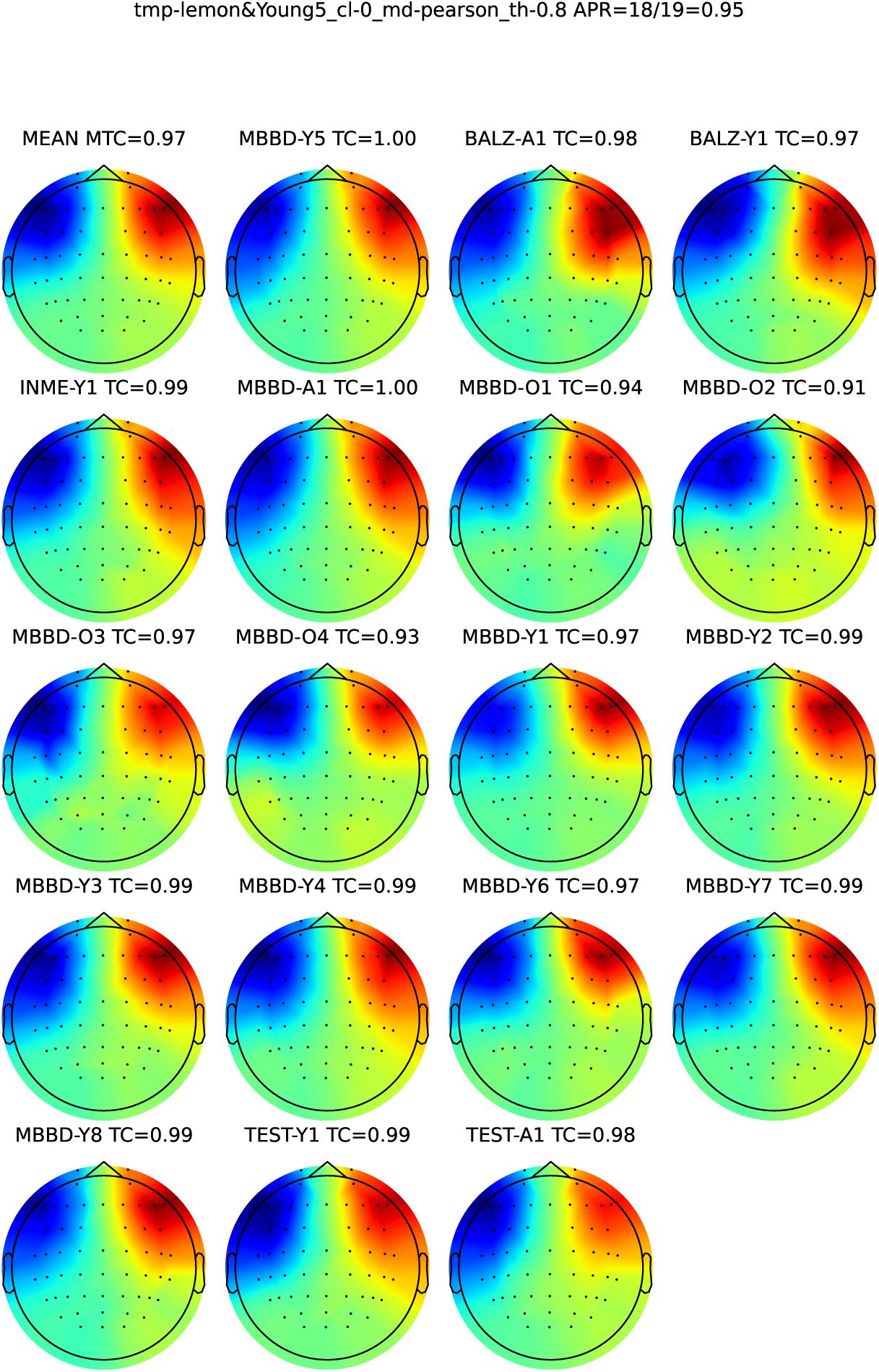
Component 1.

### D.2 Component 2

See Figure D3.

**Fig. D3.**
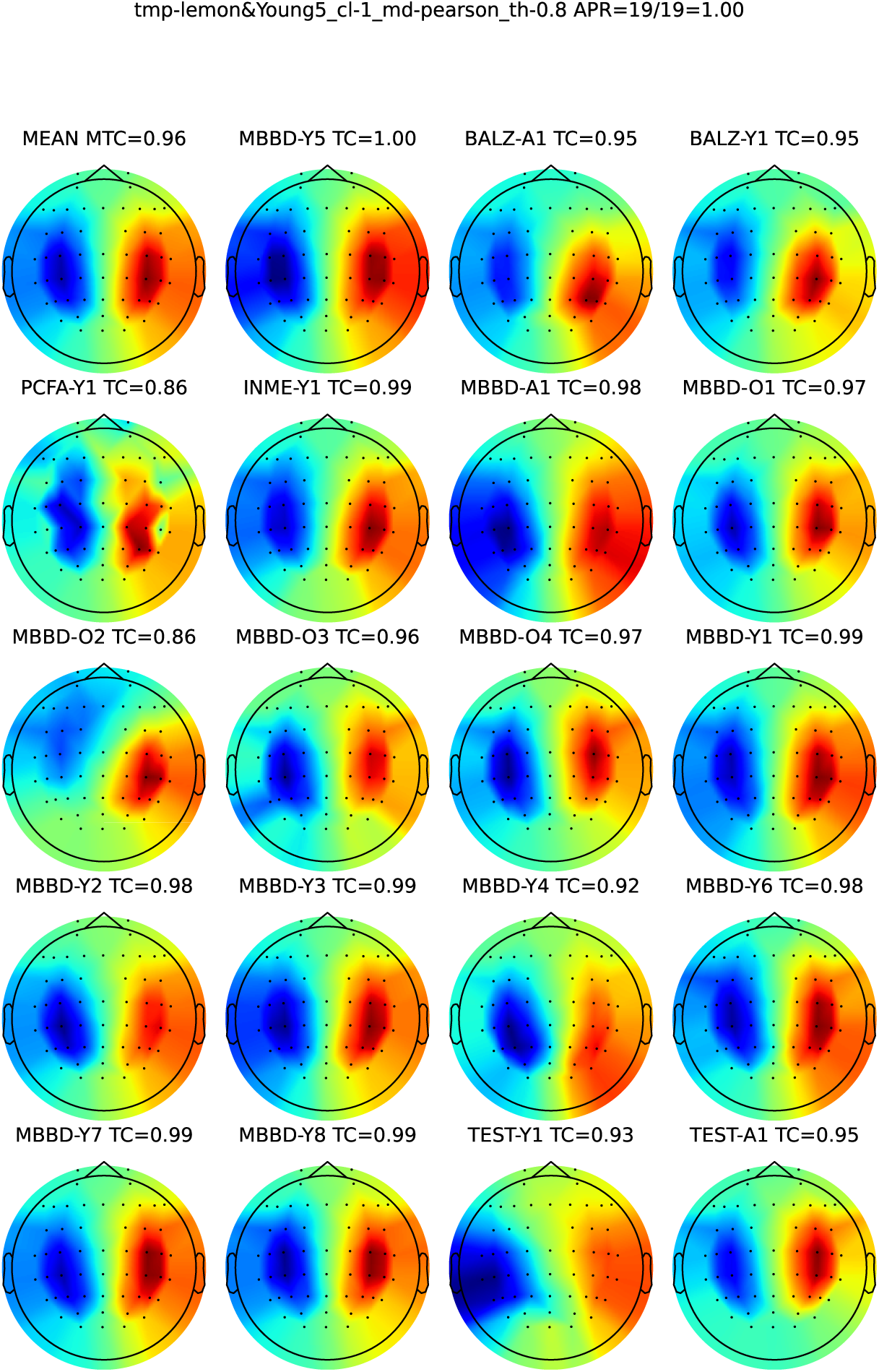
Component 2.

### D.3 Component 3

See Figure D4.

**Fig. D4.**
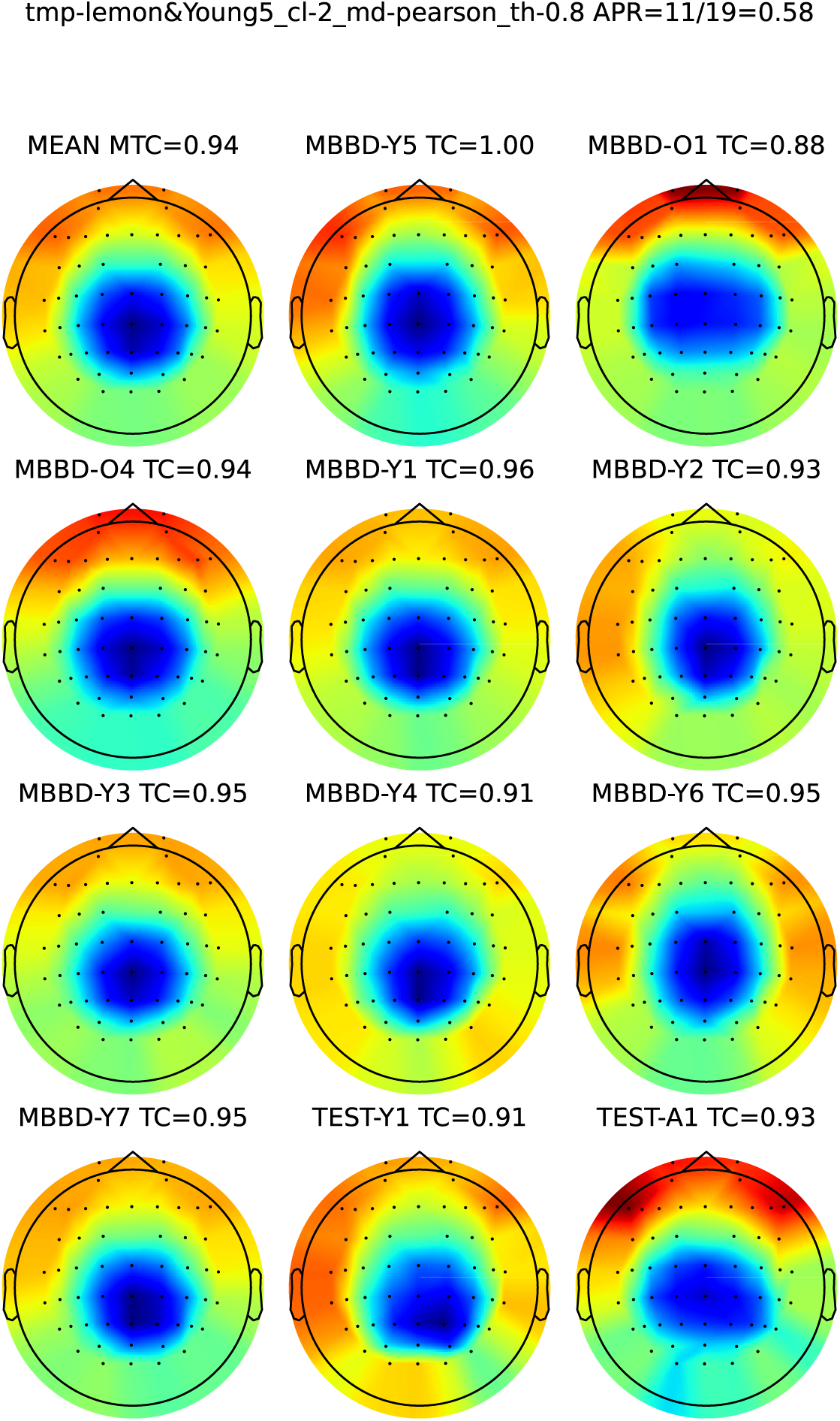
Component 3.

### D.4 Component 4

See Figure D5.

**Fig. D5.**
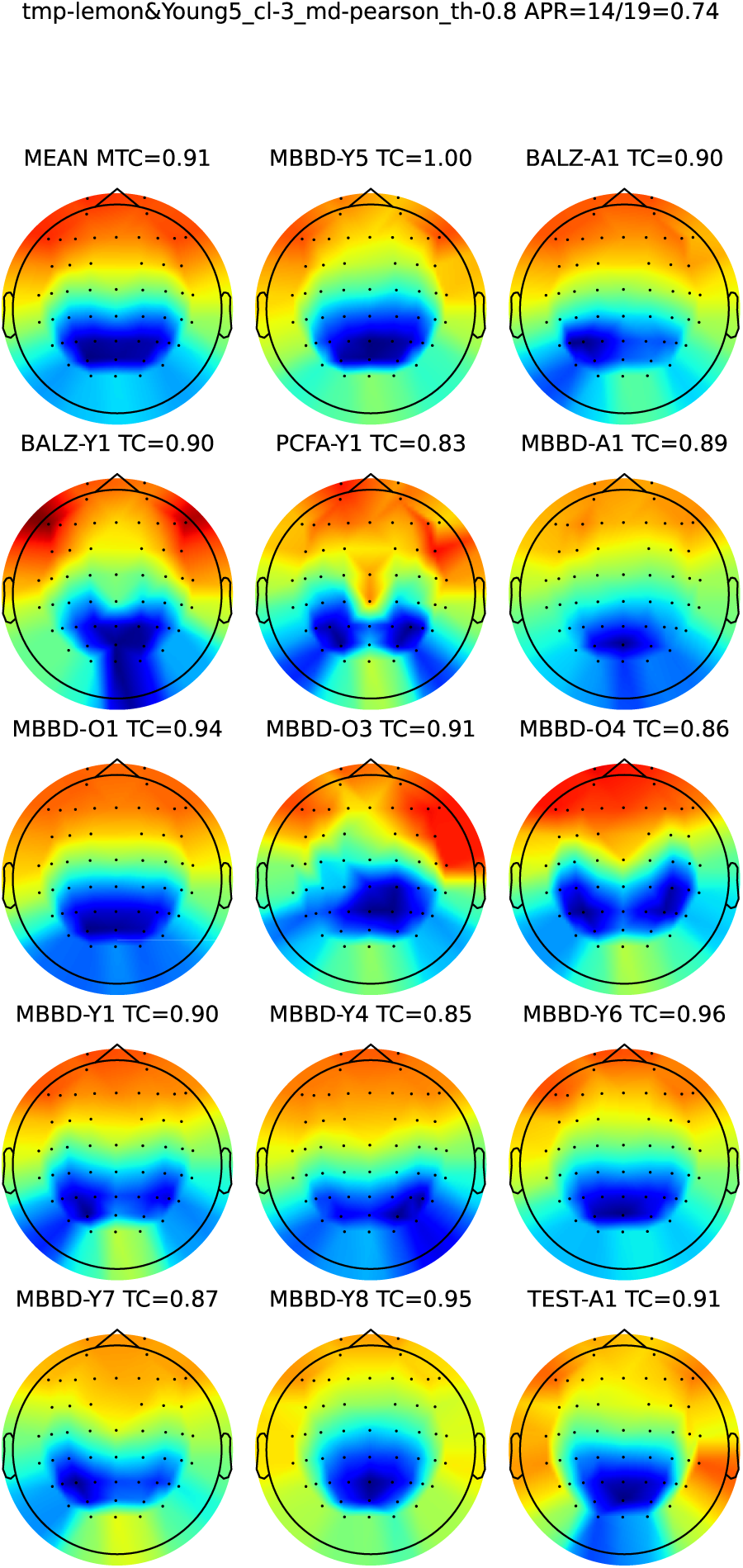
Component 4.

### D.5 Component 5

See Figure D6.

**Fig. D6.**
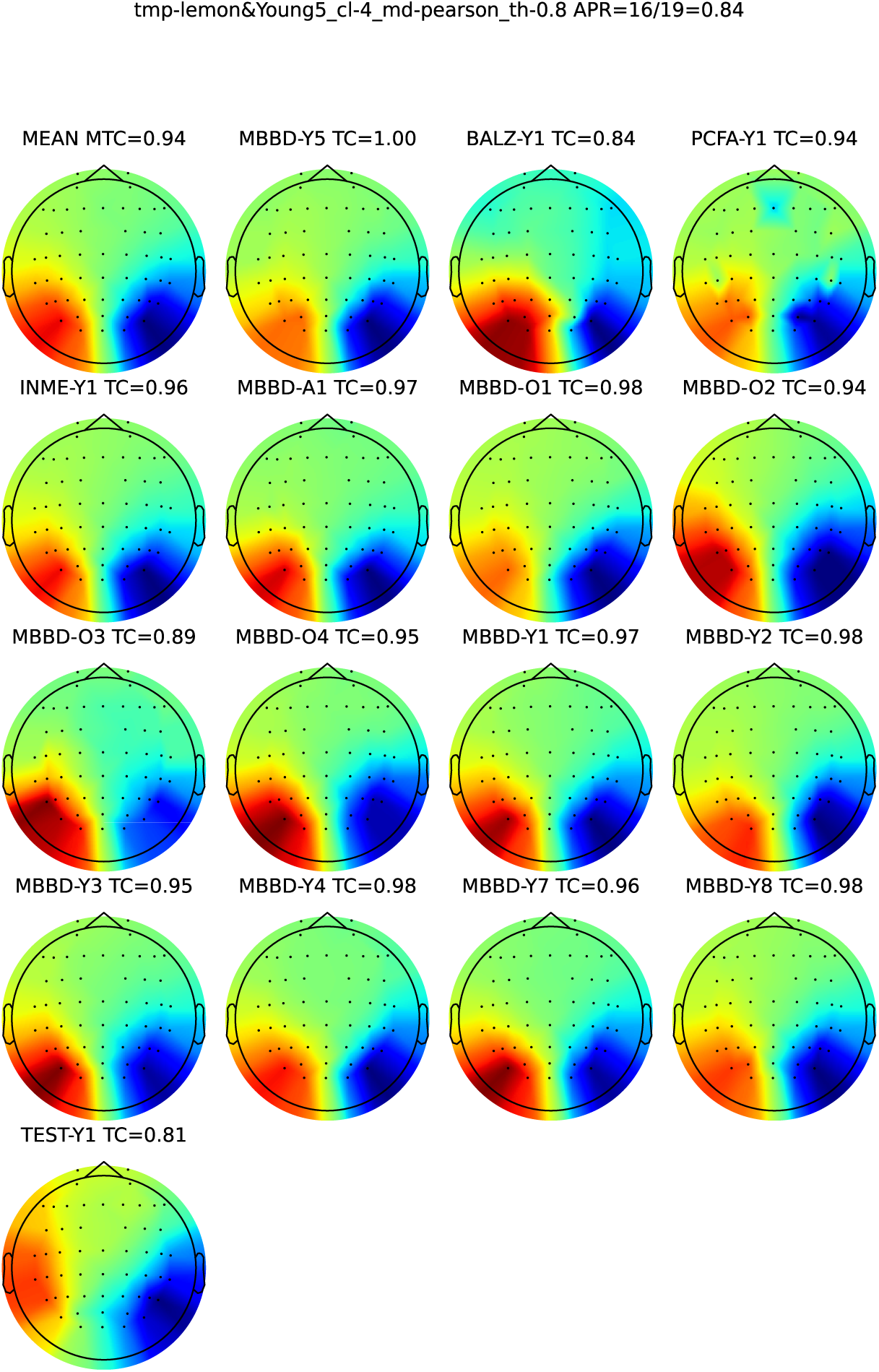
Component 5.

### D.6 Component 6

See Figure D7.

**Fig. D7.**
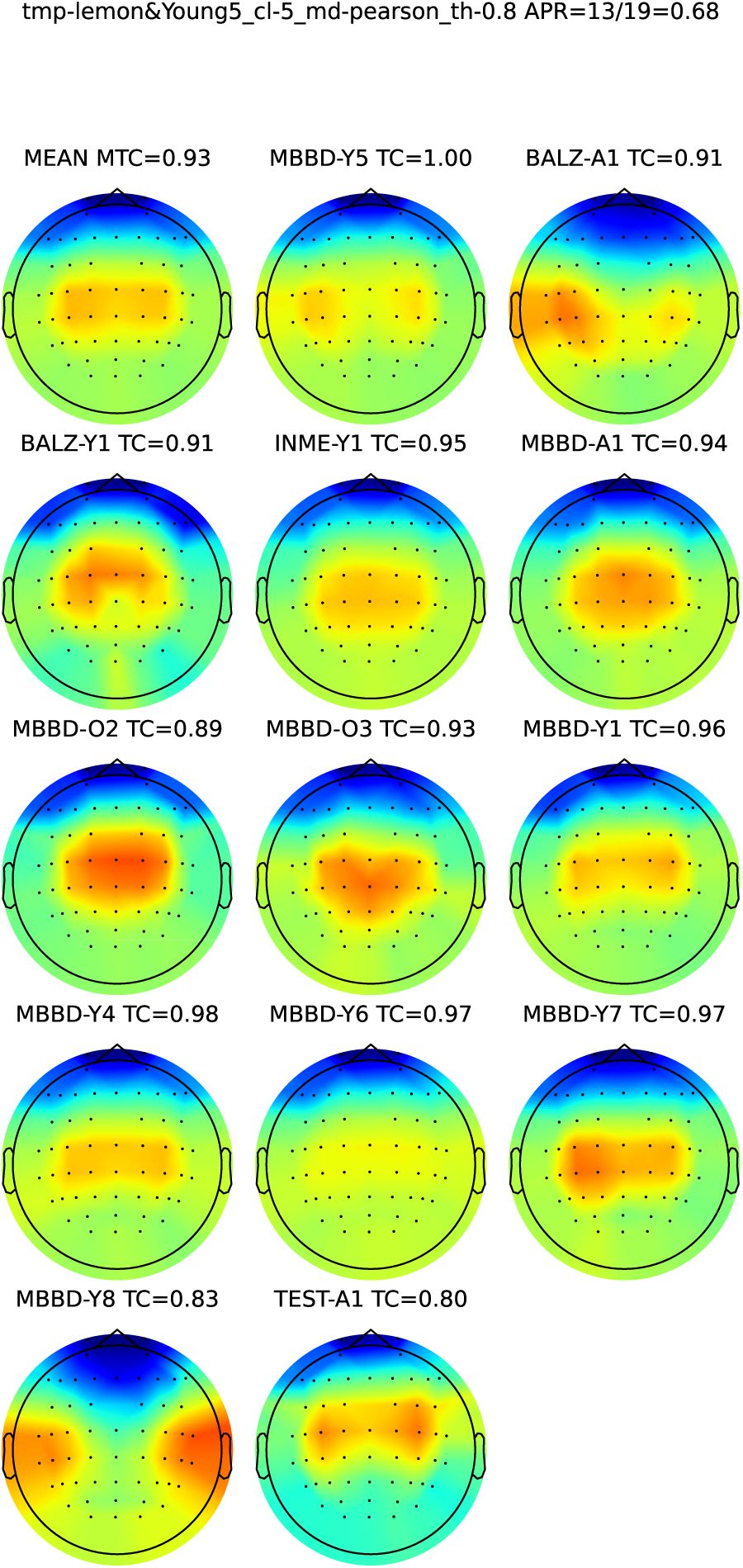
Component 6.

### D.7 Component 7

See Figure D8.

**Fig. D8.**
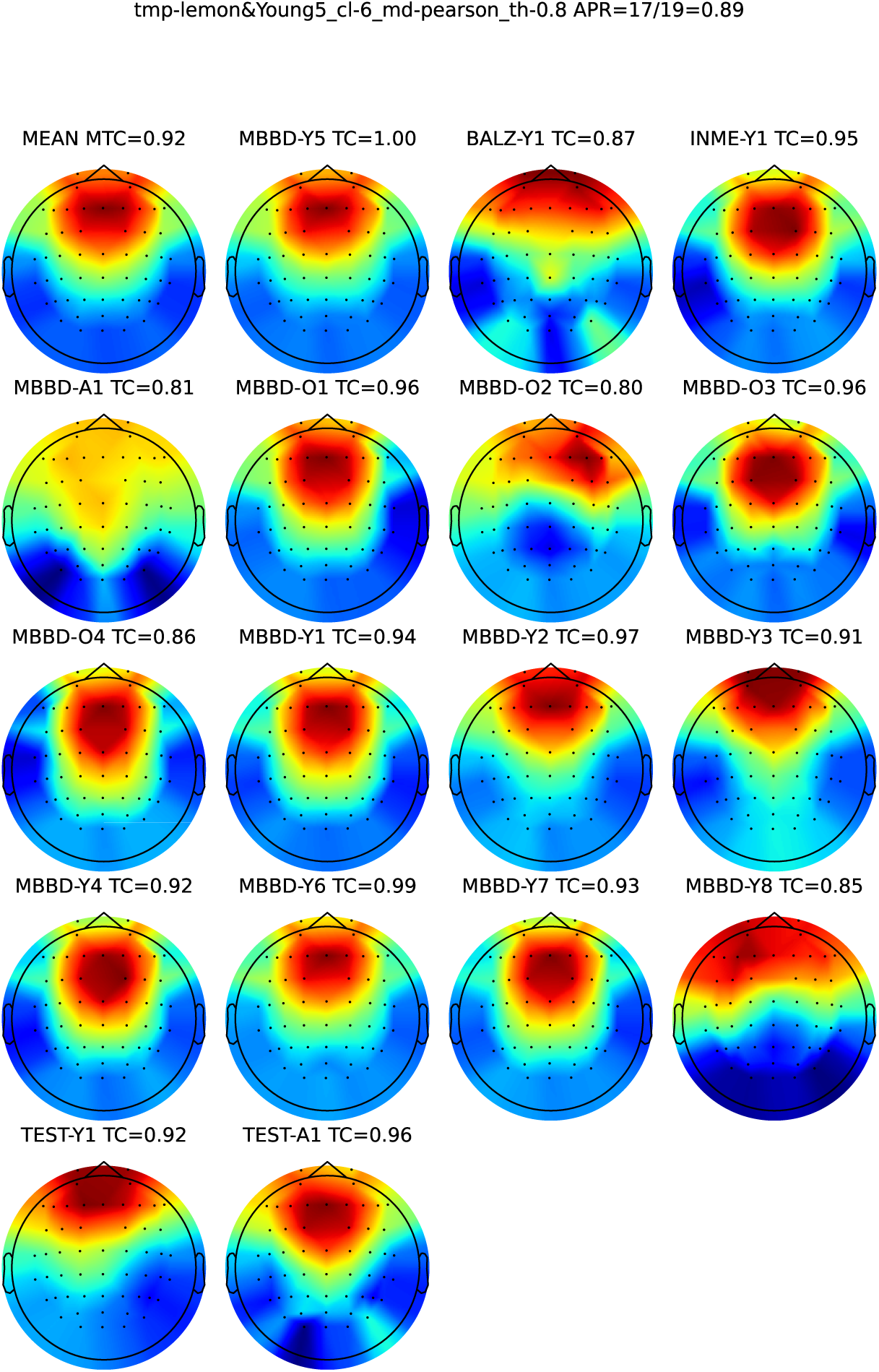
Component 7.

### D.8 Component 8

See Figure D9.

**Fig. D9.**
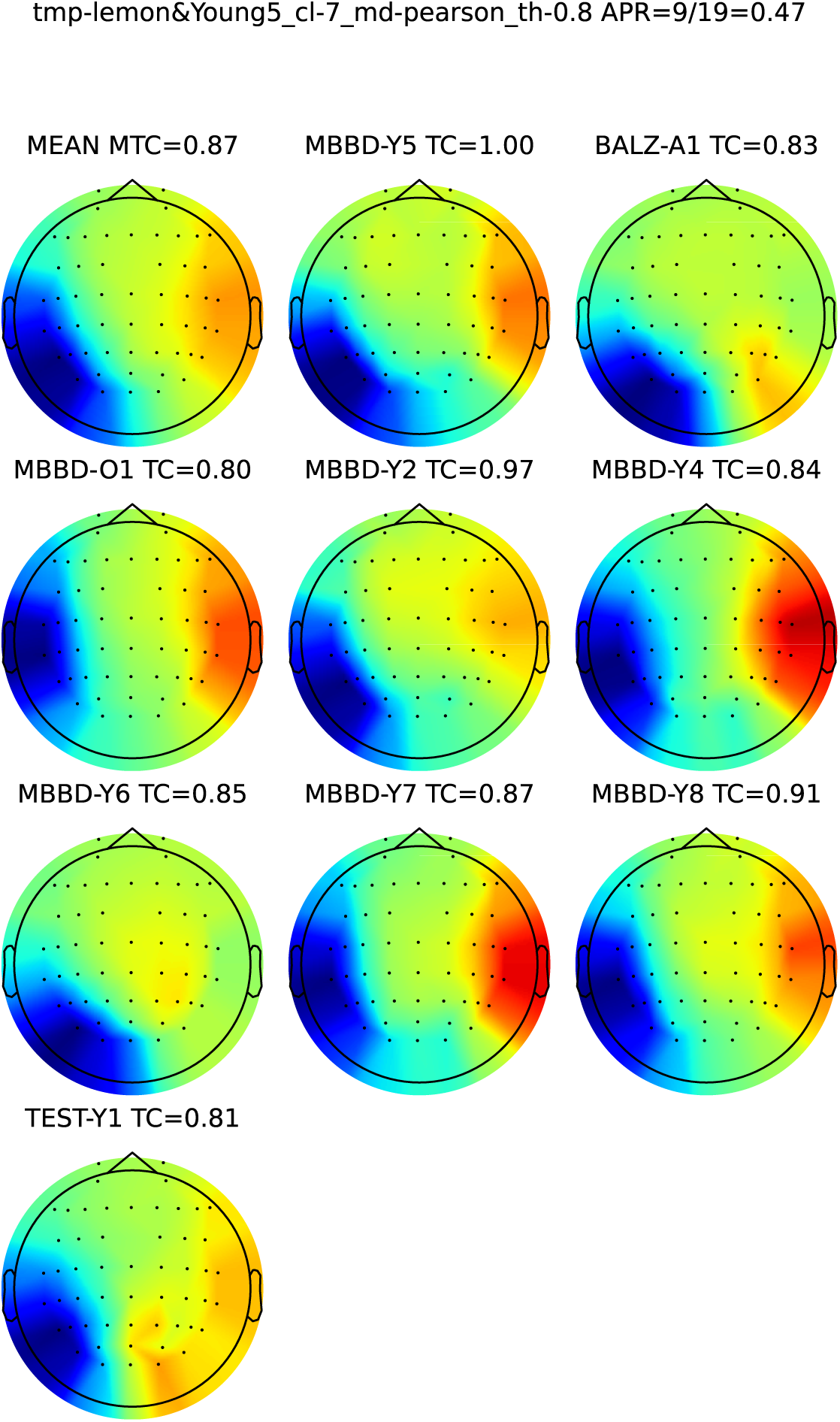
Component 8.

### D.9 Component 9

See Figure D10.

**Fig. D10.**
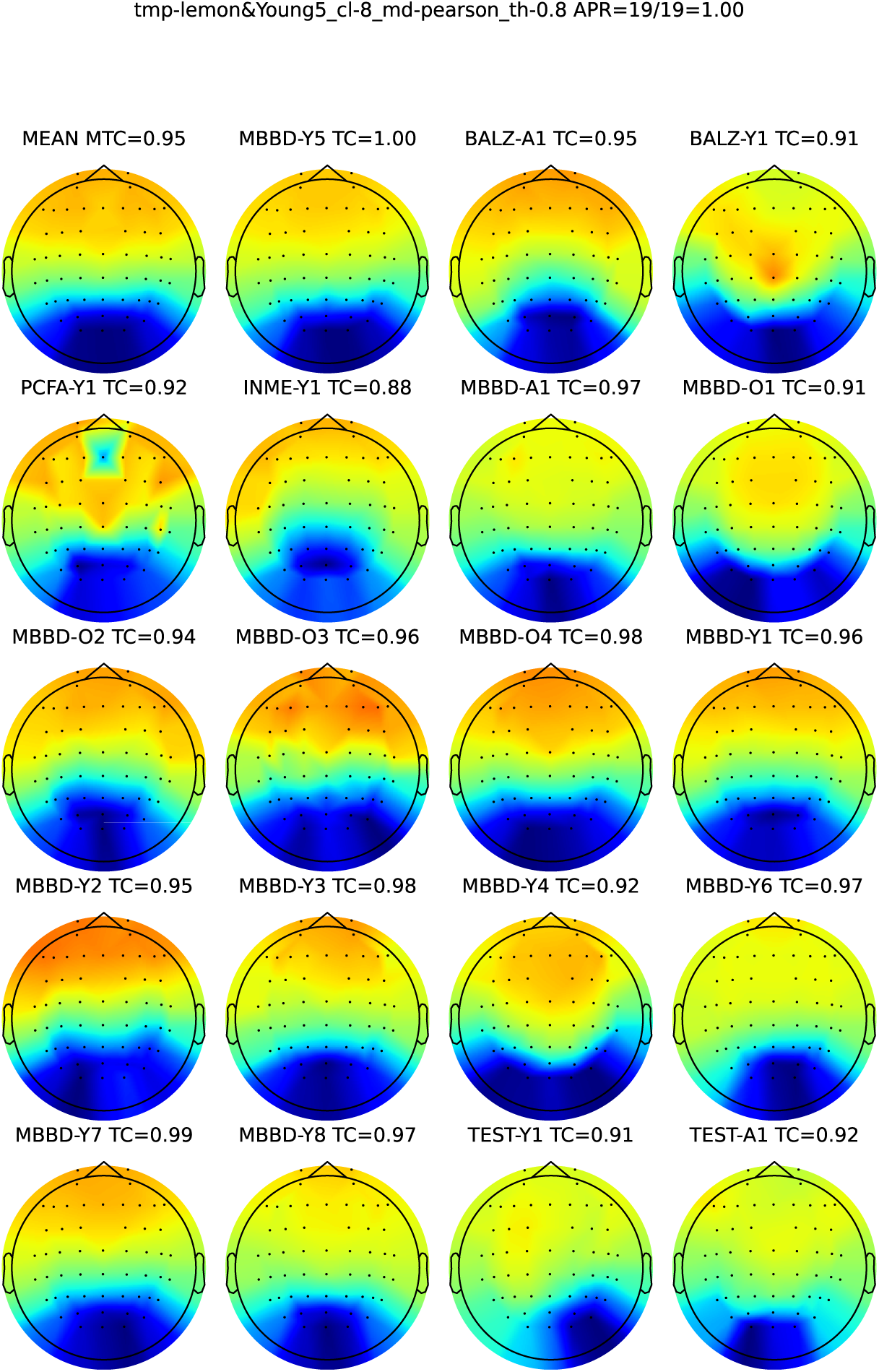
Component 9.

### D.10 Component 10

See Figure D11.

**Fig. D11.**
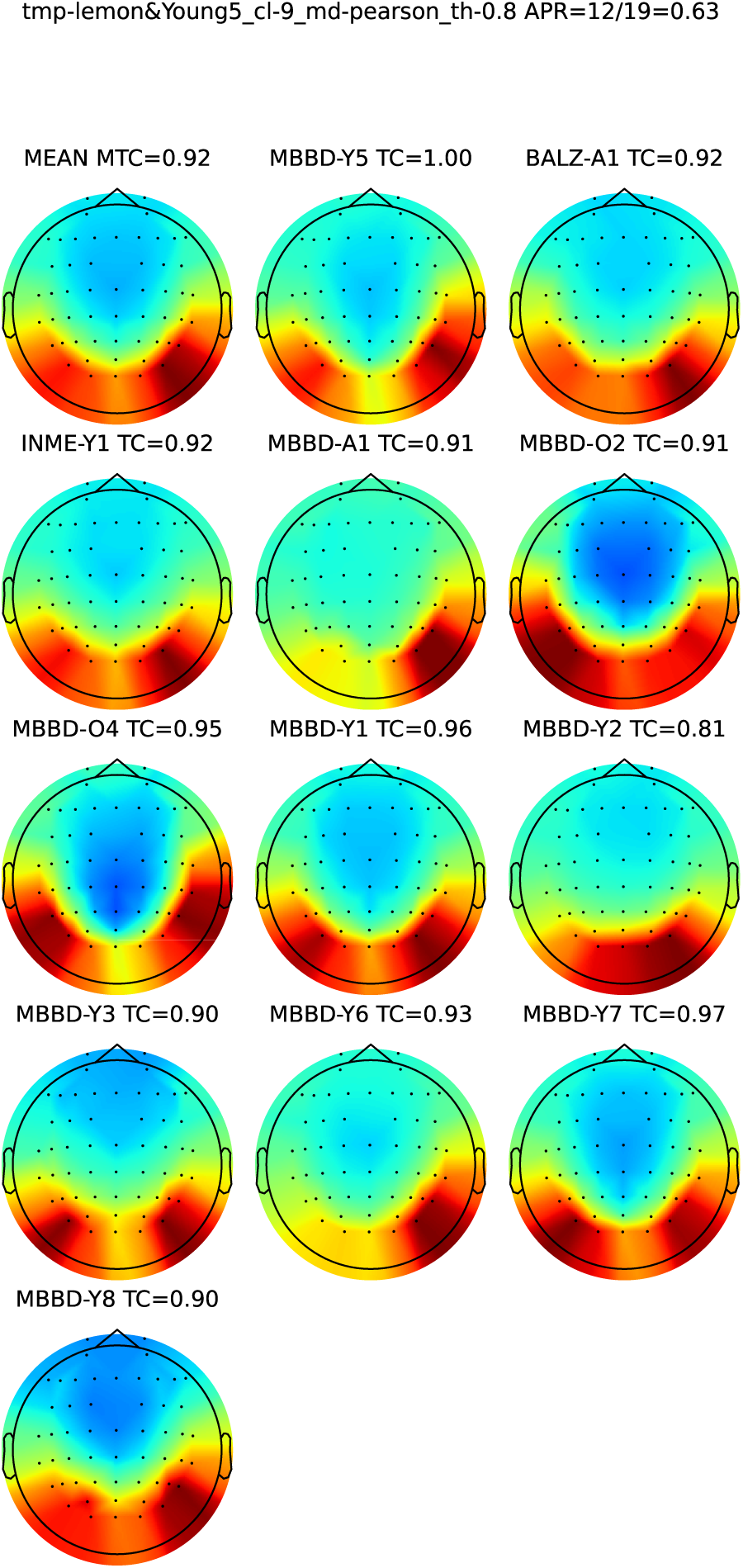
Component 10.

## Appendix E Skipping the ICASSO

To reduce computational resources, only the averaged scalp maps rather than the full set of results are included. The averaged scalp maps when clustering on all individual 285 (19 groups × 15 iterations) runs of the ICASSO can be seen in the following image (MTC: mean correlation to the template component, APR: appearance ratio). APR and MTC values have minor differences to what is presented on Figure E12. The overall trends are maintained.

**Fig. E12.**
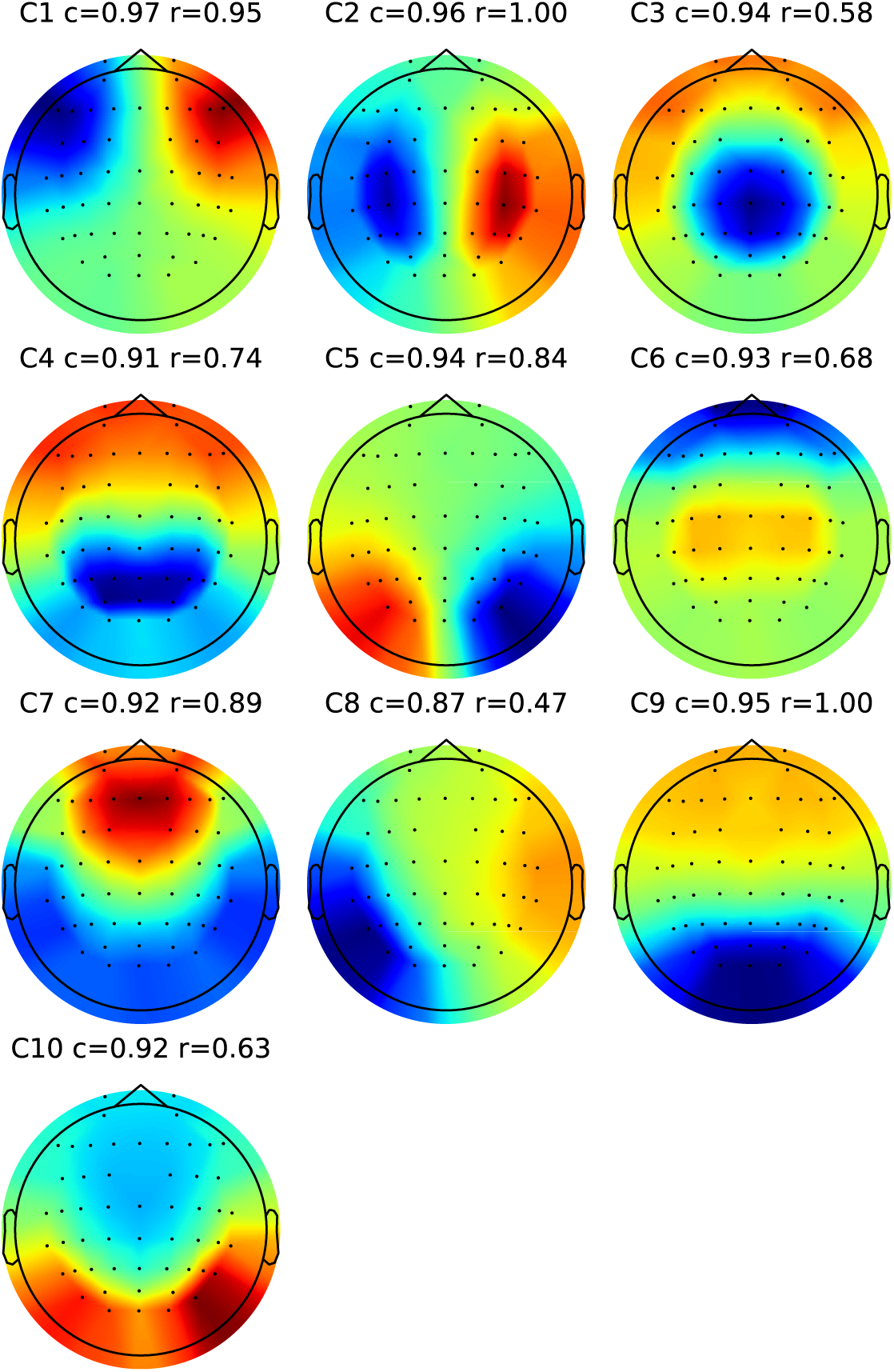
Clusters without relying on the ICASSO.

Each of these clusters have a large number of scalp maps inside, for example, for the bilateral motor component we have Figure E13:

**Fig. E13.**
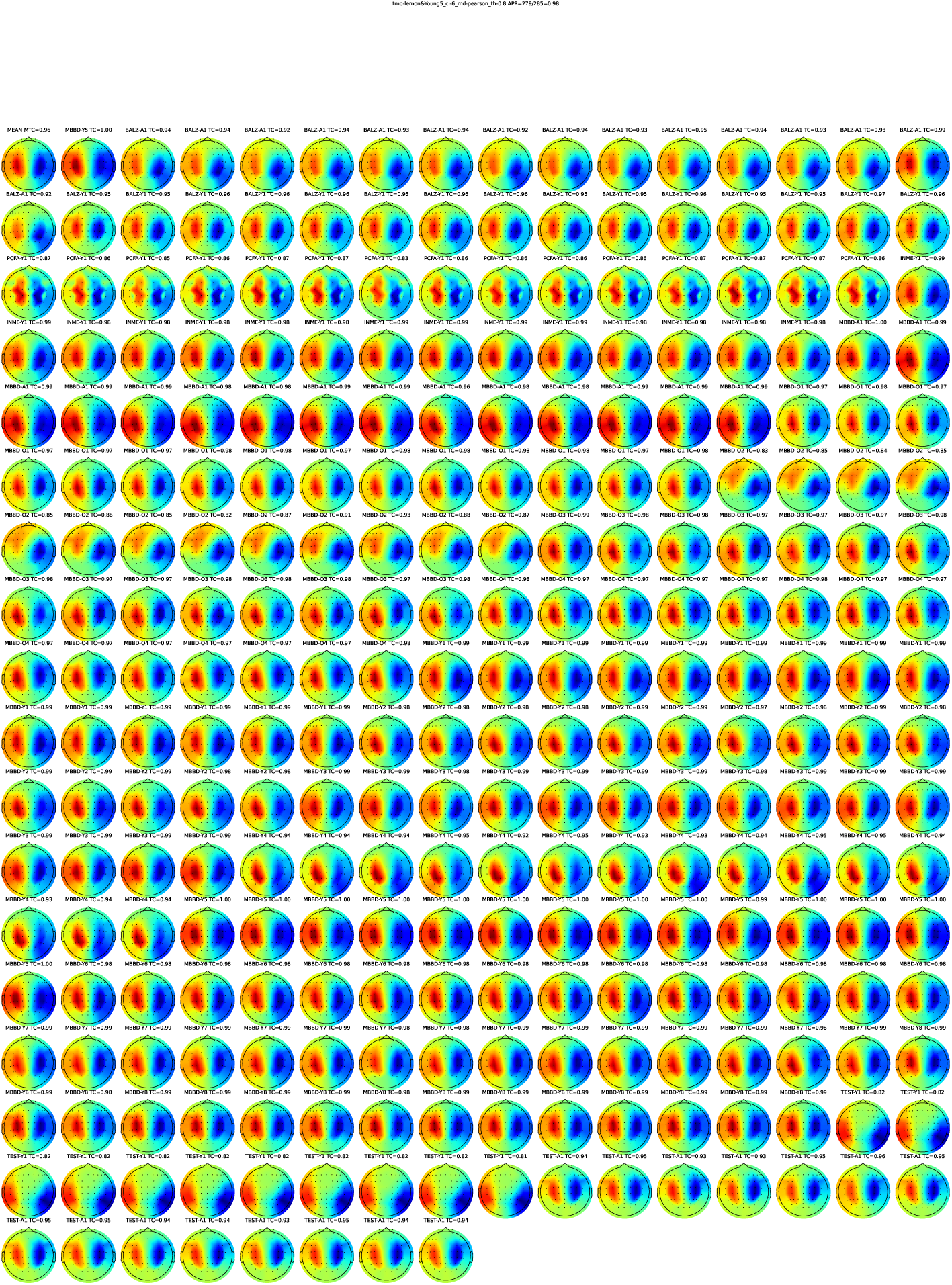
Bilateral motor cluster without the ICASSO.

## Notes

### Competing Interest Statement

The authors have declared no competing interest.

### Summary of Updates

Revised writing of the introduction. Improved discussion by including potential applications of spatial filters found through gICA and their relation to microstate maps. Corrected arrangement of the appendix.

